# Class II Contact-Dependent growth Inhibition (CDI) systems allow for broad-range cross-species toxin delivery

**DOI:** 10.1101/514810

**Authors:** Petra Virtanen, Marcus Wäneskog, Sanna Koskiniemi

## Abstract

Contact-dependent growth inhibition (CDI) allows bacteria to recognize kin cells in mixed bacterial populations. In *Escherichia coli*, CDI mediated effector delivery has been shown to be species-specific, with a preference for the own strain over others. This specificity is achieved through an interaction between a receptor-binding domain in the CdiA protein and its cognate receptor protein on the target cell. But how conserved this specificity is has not previously been investigated in detail. Here we show that three different class II CdiA receptor-binding domains and their *Enterobacter cloacae* analog are highly promiscuous, allowing for efficient effector delivery into several different *Enterobacteriaceae species*, including *Escherichia*, *Enterobacter*, *Klebsiella and Salmonella* spp. In addition, although we observe a preference for some receptors over others, this did not limit cross-species effector delivery, suggesting that class II CdiA proteins can allow for broad-range and cross-species growth inhibition in mixed bacterial populations.

## Introduction

Bacteria live in complex microbial communities where they interact and compete with other microbes for resources. A common strategy in microbial communities is to share resources and divide labor between individuals. In order for a species to benefit from resource sharing the ability to limit the use of shared goods to ones own kin is important. To that end, bacteria use a number of different strategies including toxin delivery systems for non-self-exclusion (Wall, 2016). Delivery of anti-bacterial toxins to neighboring cells can occur through different mechanisms including cell-cell contact mediated effector delivery by type IV (Souza et al., 2015), V (Aoki et al., 2005), VI (Hood et al., 2010), and VII (Z. Cao, Casabona, Kneuper, Chalmers, & Palmer, 2016) secretion systems. Common for all these toxin delivery systems is that kin-cells express an immunity or anti-toxin protein that protects against the toxic activity of its cognate effector, allowing kin cells to survive effector delivery when non-kin cells are killed or growth inhibited (Aoki et al., 2010; Aoki et al., 2005; Z. Cao et al., 2016; Hood et al., 2010; Russell, Peterson, & Mougous, 2014)

Contact-dependent growth inhibition (CDI) systems belong to the type 5 secretion system (T5SS) subclass B, also known as two-partner secretion (TPS) systems (Hayes, Aoki, & Low, 2010). During CDI, the *β*-barrel CdiB protein exports and presents the "stick-like" CdiA protein on the cell-surface (Leo, Grin, & Linke, 2012; Ruhe, Low, & Hayes, 2013; Ruhe et al., 2018). The delivered toxic effector is encoded in the C-terminal domain of the CdiA protein (CdiA-CT). Upon interaction with a receptor on the cell surface of a neighboring cell, the C-terminal domain is cleaved off and delivered into the recipient cell (Aoki et al., 2010; Ruhe et al., 2018; Webb et al., 2013). The toxic activities of the C-terminal encoded toxic effectors range from nucleases that degrade tRNA, rRNA and DNA to ionophore toxins that dissipate the proton motive force of the bacterial cell (Aoki et al., 2010; Aoki, Webb, Braaten, & Low, 2009; Morse et al., 2012). To protect themselves from auto-inhibition bacteria with CdiA toxins express a small CdiI immunity protein, which binds and forms a tight complex with its cognate toxin, neutralizing its toxic activity (Aoki et al., 2005; Morse et al., 2012). *E. coli* CdiA proteins can be divided into different classes (Class I-V) based on sequence homology. The sequence variation between Class I, II and III CdiA proteins is mainly found in the receptor-binding domain (RBD), with the exception of the toxic C-terminal domains (which are highly variable as they encode for toxins with diverse toxic activities). In the first identified Class I CdiA protein of *E. coli* EC93 the RBD is found in the middle of the CdiA protein (residues ~1300-1600aa and ~1900-2300aa) (Ruhe et al., 2017). These CdiA RBD interact with different outer-membrane proteins on the surface of targeted cells; BamA for Class I (Aoki et al., 2008; Ruhe, Wallace, Low, & Hayes, 2013), OmpC-OmpF for Class II (Beck et al., 2016) and Stx for Class III (Ruhe et al., 2017), enabling delivery of effectors to the recipient cell. Previous studies suggest that effector delivery by class I CdiA proteins is strictly species-specific and limited to *E. coli* (Ruhe, Wallace, et al., 2013) and that the class II CdiA proteins are strain-specific and able to discriminate between different strains of *E. coli*, with a preference for the “own” strain over others (Beck et al., 2016). This specificity is achieved by differences in the protein sequences of the extracellular loops of the receptors, which presumably affects the binding affinity between the receptor and the RBD of CdiA. After being delivered through the outer membrane of a target cell, the C-terminal toxin domains of many CdiA toxins associate with different inner-membrane proteins for translocation into or through the inner-membrane (depending on the activity of the toxin). The latter interaction does not seem species-specific as toxins from many different species can be attached to the CdiA-stick and be efficiently delivered into *E. coli* cells (Willett, Gucinski, Fatherree, Low, & Hayes, 2015).

We were interested in finding out more about the specificity of binding between class II CdiA RBD and their cognate receptors, hoping to learn more about the role that CDI systems play in kin-recognition. Previous studies on kin recognition proteins like TraA in *Myxococus xanthus*, show that single amino acid changes are sufficient for differential binding between proteins and their cognate receptors (P. Cao & Wall, 2017) and we wanted to investigate if this is also the case for the interaction between CdiA and the OmpC component of the receptor, whose extracellular loops were previously been shown to drive specificity (Beck et al., 2016). We used bioinformatics to find similar but not identical CdiA RBD toinvestigate if small changes in the RBD changed the receptor recognition and thus the species-specificity of CdiA toxin delivery. Somewhat surprisingly, we find that three closely homologous *E. coli* class II CdiA RBD allow for delivery of toxic effectors into many different *Enterobacteriaceae* spp. including; *Enterobacter cloacae* and *aerogenes, Klebsiella pneumoniae* and *Salmonella typhimurium*, suggesting that class II CDI is a broad-range inter-species competition system. Additionally, our results suggest that although receptor binding correlates well with inhibition during continuous agitation, strong receptor binding is not required for CDI on solid media. Taken together, our results suggest that class II CdiA molecules allow for kin recognition through inhibition of other bacteria and at the same time illustrate how the versatility of CDI as a competition and kin recognition system changes, depending on the environmental context.

## Results

### Identification of CdiA proteins in *Enterobacteriaceae* spp

To increase our understanding of the interaction between the class II CdiA RBD and its cognate OmpC receptor, we set out to bioinformatically identify similar but not identical CdiA receptor-binding domains with the intention of studying if small changes in the receptor-binding domain affects receptor-specificity. Our bioinformatic analyses identified identical and closely homologous class II CdiA receptor-binding domains in *E. coli* strains with quite different OmpC protein sequences. For example, *E. coli* UPEC F11 has an identical CdiA binding domain to that found in the previously studied UPEC 536 (Fig. S1) but has very different extracellular loops of OmpC (Fig. S2). In addition, closely homologous binding domains (few aa differences) to the CdiA protein of *E. coli* UPEC F11 (CdiA^F11^) were identified in *Salmonella typhi, E. coli* CFT073/*E. coli* Nissle 1917 (Fig. S1), which also have significantly different OmpC extracellular loops from UPEC 536, UPEC F11 and each other (with the exception of *E. coli* CFT073 and Nissle 1917 where both binding domains and OmpC sequences were identical) (Fig. S2). Thus, these findings suggest that species-specificity could be achieved by very small amino acid differences in the receptor and/or receptor-binding domain.

### Class II CdiA-OmpC dependent effector delivery is promiscuous

To test how the differences between OmpC proteins affected class II mediated toxin delivery, we replaced the chromosomal *ompC* ORF of *E. coli* MG1655 with the *ompC* from *E. coli* strains UPEC F11 or Nissle 1917/CFT073, as well as the *ompC* from *Enterobacter cloacae* and *Salmonella typhimurium/typhi* (OmpC’s from *S. typhimurium* and *typhi* are identical). Next, we competed these strains with an *E. coli* MG1655 strain expressing a chimeric CdiA protein with the receptor-binding domain from UPEC F11 (CdiA^F11^) from a medium copy (ColE1) plasmid containing the *cdiBAI* locus from EC93 under its native promoter. Surprisingly, cells expressing CdiA^F11^ outcompeted all target strains during co-cultivation in liquid LB media (Fig. 1A, dark green bars). Inhibition varied between 4-logs for the *E. coli* UPEC F11 and *E. cloacae* OmpC’s (OmpC^F11^ and OmpC^ECL^ respectively) to 1-log for the OmpC of CFT073 (OmpC^CFT073^) (Fig. 1A, dark green bars). Cells expressing OmpC variants from *S. typhimurium/typhi* (OmpC^*Sty*^), or the native OmpC of *E. coli* MG1655 (OmpC^K12^) were outcompeted with 2-logs (Fig. 1A, dark green bars). Furthermore, cells expressing CdiA^F11^ were not able to outcompete cells expressing CdiI immunity protein irrespective of their OmpC, suggesting that the observed ability to outcompete was indeed mediated by toxic effector delivery into the different strains (Fig. 1A, light green bars). To further confirm that the observed growth inhibition was due to toxin delivery, we used cells lacking the *ompC* gene (Δ*ompC).* MG1655 cells lacking *ompC* were not outcompeted by cells expressing CdiA^F11^ (Fig. 1A), further confirming that the observed inhibition was mediated by CDI and that OmpC indeed functions as a receptor for CdiA^F11^. Notably, expression of *cdiI* came with a similar fitness cost for the cells as expressing *cdiBAI*.

**Figure 1.**
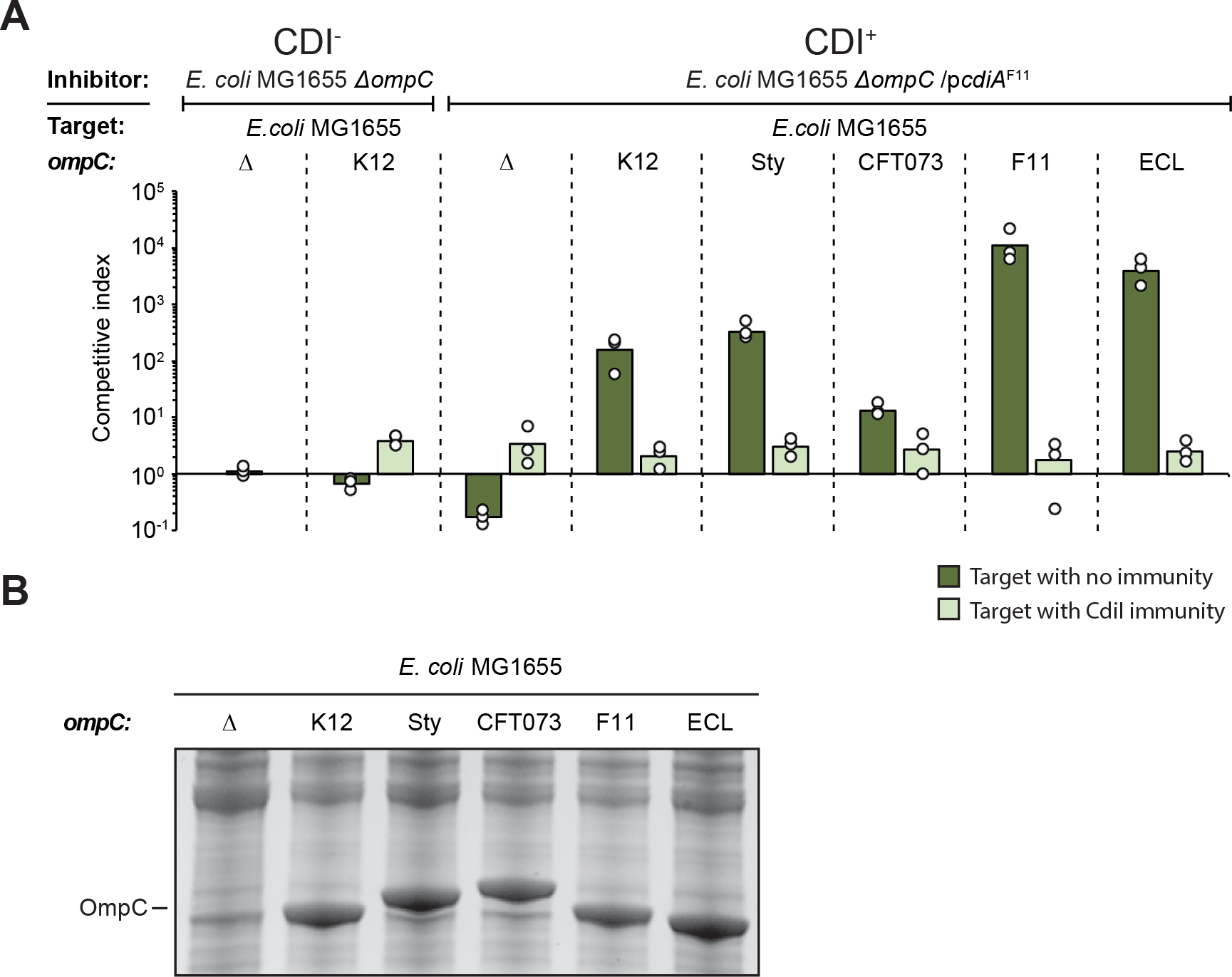
Class II CdiA proteins are able to deliver effectors to cells expressing the OmpC receptor from other strains and species. **A)** Average competitive index of cells expressing CdiA^F11^ after co-culturing with MG1655 cells expressing different OmpC’s from the native MG1655 *ompC* locus with (light green bars) or without (dark green bars) CdiI expressed from plasmid (n=3 biological replicates). Cells were co-cultured for 5h in liquid LB media. Individual data points of the biological replicates are shown as black and white circles. **B)** MG1655 cells expressing different OmpC’s from the native MG1655 *ompC* locus from Fig. 1A were grown in LB and outer-membrane fractions were enriched and separated by SDS-PAGE.

We did observe differences in the level of inhibition between cells expressing different OmpC variants. There are two possible explanations to this observation, i) some OmpC’s allow more efficient toxin delivery or ii) the target strains could express different levels of the OmpC proteins. To test the latter we used SYPRO Ruby (Thermo Scientific) protein staining to measure OmpC levels in the different strains. OmpC protein levels were very similar in the different target strains (Fig. 1B), suggesting that OmpC levels are not the reason for the difference in inhibition between these strains. Taken together, our results suggest that class II CdiA mediated effector delivery is not species-specific, but that there could be a difference in class II CdiA receptor specificity.

### OmpC levels are important for*cdiBAI* mediated growth inhibition

Our findings of class II CdiA cross-species growth inhibition are in contrast with previous findings where cells expressing a class II CdiA protein from UPEC 536 (CdiA^UPEC^) were not able to deliver effectors to *E. coli* cells expressing OmpC from *S. typhimurium* (Beck et al., 2016). The RBD of UPEC 536 and F11 are identical, but to our surprise, MG1655 strains expressing OmpC^*Sty*^ were inhibited as efficiently as wild type MG1655 cells (OmpC^K12^) by inhibitor cells expressing CdiA^F11^. In the previous study, a plasmid based construct was used to express OmpC^*Sty*^ from a leaky pTac promoter (Beck et al., 2016). Our construct has the *ompC* ORF from *S. typhimurium/typhi* inserted on the chromosome under the native *ompC* promoter and should, under these conditions, express roughly 100,000 OmpC molecules/cell (Schuman, 2006). Thus, an obvious difference between these constructs is the expression level of OmpC. To test if OmpC expression levels are important for CdiA cross-species effector delivery, we cloned all the tested *ompC* ORFs onto a low-copy (pSC101) plasmid backbone, to be expressed from a synthetic, medium strong, constitutive promoter; PJ23101 (Kelly et al., 2009). Cells expressing CdiA^F11^ were able to outcompete MG1655 cells expressing low levels of OmpC^F11^ or OmpC^ECL^ in liquid LB media, whereas target cells expressing any of the other OmpC variants were not inhibited (Fig. S3A). This is different from the growth inhibition observed of strains expressing the chromosomally encoded OmpC’s, where all target cells, regardless of OmpC variant, were inhibited (Fig. 1A). These results are in line with the previously reported lack of inhibition of target cells expressing OmpC^*Sty*^ (Beck et al., 2016). We confirmed that our plasmid constructs expressed lower levels of OmpC than our chromosomal constructs using SDS-PAGE (Fig. S3B) and western blot (Fig. S3C). The chromosomal constructs expressed ~550-fold more OmpC than the plasmid constructs during growth in liquid LB media (Fig. S3C). Thus, the level of the OmpC receptor does affect effector delivery, which could be the reason why a broad-range cross-species effector delivery has not previously been observed.

### Inhibition correlates with CdiA-receptor binding

Our results suggest that cells expressing CdiA^F11^ proteins are able to inhibit the growth of cells expressing OmpC from F11 and *E. cloacae* more than cells expressing other OmpC variants. This does not necessarily mean that the binding interactions between the CdiA and the different OmpC proteins vary. To test if CdiA proteins with the class II binding domain have different binding affinity for OmpC’s from different species, we used a previously described cell-cell binding assay (Aoki et al., 2008). In short; inhibitor cells expressing CdiA^F11^ were modified to constitutively express one fluorophore (sYFP2) and target cells with different OmpC variants (OmpC^K12^, OmpC^Sty^, OmpC^CFT073^, OmpC^F11^ or OmpC^ECL^) expressing another fluorophore (dTomato). Inhibitor and target cells were mixed and after 40 min of co-cultivation at a high cell-density in liquid LB with shaking the fraction of dTomato+ target cells (with different OmpC variants) bound to sYFP2^+^ inhibitor cells (with CdiA^F11^) were analyzed by flow cytometry (Fig. 2A). To control for receptor independent binding target cells lacking the OmpC receptor were used. Approximately 34% of target cells expressing OmpC^F11^ were bound to cells expressing CdiA^F11^, compared to 26% of cells expressing OmpC^ECL^, 15% of cells expressing OmpC^K12^ and 12% of cells expressing OmpC^CFT^ (Fig. 2A-B). Around 10% of *ΔompC* target cells were bound to inhibitors (receptor independent cell-cell interactions) (Fig. 2B). For target cells expressing OmpC^Sty^, binding above background levels (10%) could not be detected, even though these cells were inhibited to the same extent as cells expressing OmpC^K12^ (Fig. 1A).

**Figure 2.**
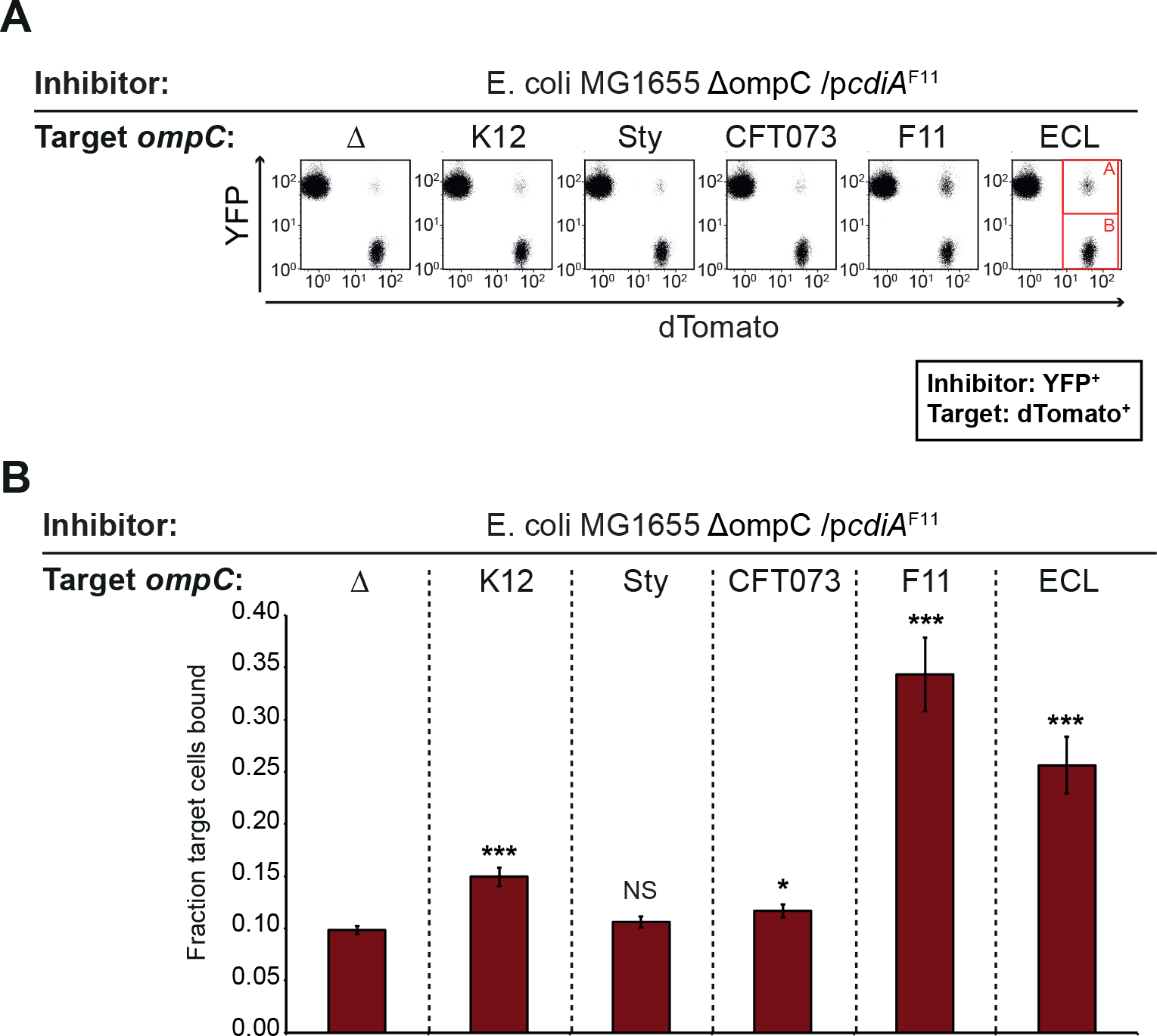
CdiA-OmpC mediated cell-cell binding. YFP+ MG1655 cells expressing CdiA^F11^ were mixed with dTomato+ MG1655 cells expressing none (*ΔompC*) or different OmpC’s from the native *ompC* locus on the MG1655 chromosome. Cell-cell binding was analyzed by flow cytometry. **A)** Representative images of flow cytometry cell-cell binding assay. Bound dTomato+ target cells (A, red square) were divided by total number of dTomato+ cells (A+B, red squares) to calculate fraction of bound target cells. **B)** Average fraction of MG1655 target cells bound to CdiA^F11^ expressing inhibitor cells (n=6 biological replicates). Error-bars are SEM. Statistical significance was determined using a two-tailed, unpaired students t-test where * = P<0.05, ** = P<0.005 and ***= P<0.0005.

We correlated relative cell-cell binding versus CDI mediated growth inhibition and could observe that there was an almost perfect polynomial correlation (P<0.01) between cell-cell binding and growth inhibition (Fig. S4). Thus, we concluded that strong cell-cell binding is not always beneficial for CDI mediated growth inhibition. As our binding assay measures steady-state receptor binding both CdiA-OmpC association and dissociation are measured in our assay (Fig. 2A-B). Thus, our results could be explained through a general crowding effect where a medium CdiA-OmpC binding-affinity results in a sequestering of inhibitor cells, allowing the dissociation rate to determine how many target cells can be inhibited (i.e. the first target cell must be released before a second target cell can be bound and inhibited). At lower steady-state binding, the dissociation rate is most likely higher, allowing more cells to be inhibited in the same timeframe. In contrast, high-affinity binding increases the likelihood of inhibitor–target cell interactions and would possible allow for cells to form larger aggregates, both of which would reduce the negative effects of a slow dissociation rate on growth inhibition. This relationship between cell-cell binding and growth inhibition is well described by a polynomial distribution and thus we concluded that the measured CdiA-OmpC interaction does correlate with observed growth inhibition in this context.

### Growth on solid media overcomes weak CdiA-OmpC binding

In their natural environment, most bacteria grow on solid surfaces or within biofilms. In such conditions, the binding affinity between CdiA and its cognate receptor might not play a significant role, as binding is not required to keep proximity to the neighboring cell. To test this, we competed MG1655 cells expressing CdiA^F11^ against MG1655 cells with low (plasmid) or high (chromosomal) expression of the different OmpC variants on solid rich defined M9 media supplemented with glucose and casamino acids (M9Glu) (for details regarding the media see the materials section). MG1655 cells expressing high levels of the different OmpC variants were inhibited similarly; around 2-log for all OmpC variants except OmpC^*Sty*^ where only 1-log of inhibition could be seen (when the fitness cost of the antitoxin expressing plasmid was taken into account) (Fig. 3A). Similarly, MG1655 cells expressing low levels of any of the OmpC variants except OmpC^*Sty*^, were also outcompeted with 2- to 3-logs on solid media, (Fig. 3B). Notably the fitness cost of carrying the CDI system (as observed by the CdiA^F11^ vs. *ΔompC* competitions) was almost abolished by the presence of the pSC101 OmpC expression plasmid in the target strains (Fig. 3B), whereas the same cost added more than 1-log of difference to the strains lacking the pSC101 OmpC expression plasmid (Δ*ompC*, Fig. 3A). Taken this into account, there was not much difference in the level of inhibition between strains expressing high or low levels of the receptor when competed on solid media (Fig. 3A-B), suggesting that neither low receptor abundance nor weak CdiA-OmpC binding affinity prevent or limit growth inhibition on solid media.

**Figure 3.**
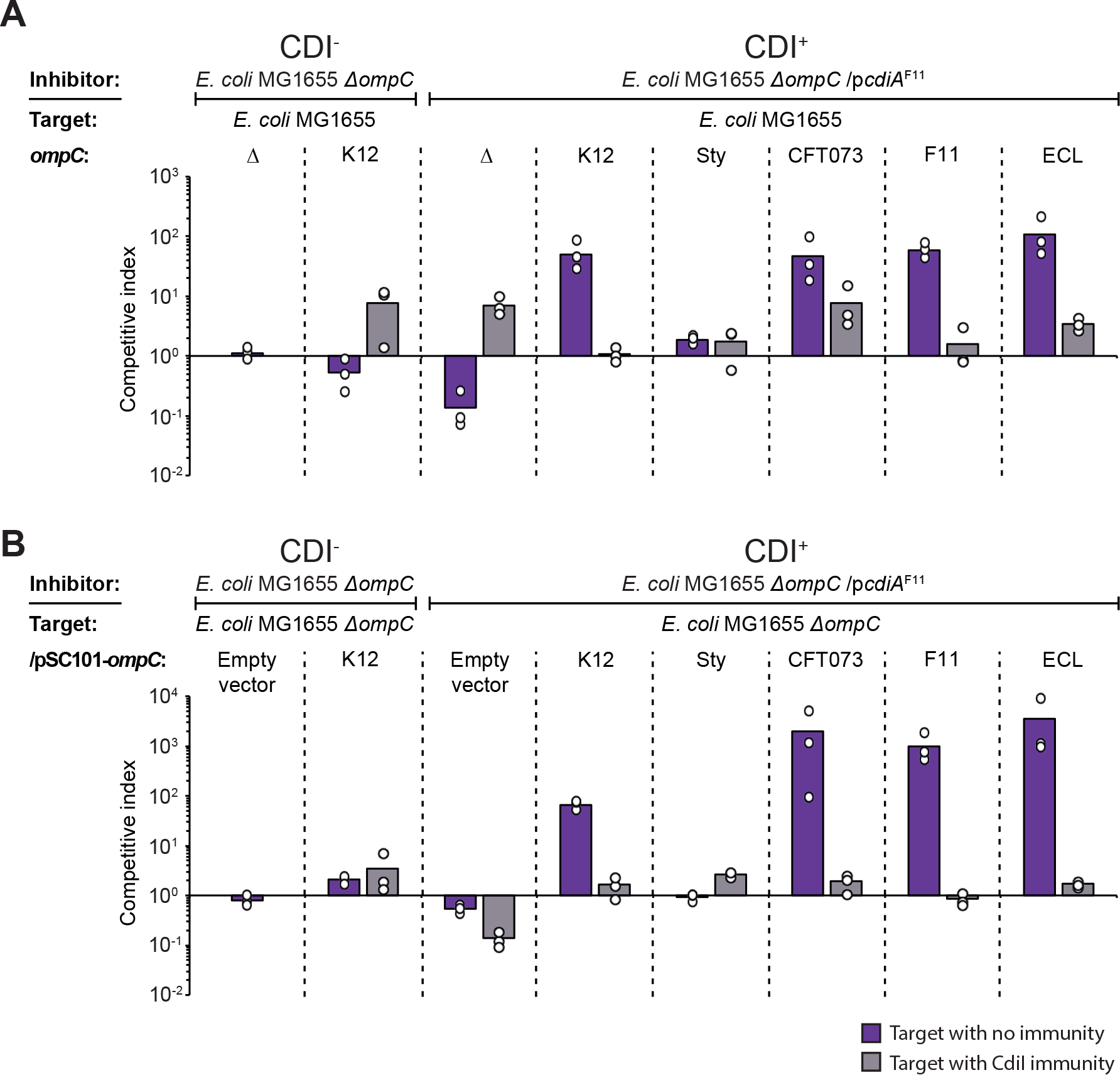
Growth on solid media overcomes weak CdiA-OmpC binding. Average competitive index of cells expressing CdiA^F11^ after co-culturing with MG1655 cells expressing different OmpC’s from the native MG1655 *ompC* locus **(A)** or from the low-copy (pSC101) plasmid **(B)** with (grey bars) or without (purple bars) CdiI expressed from plasmid (n=3 biological replicates). Cells were co-cultured for 24h on solid M9Glu media. Individual data points of the biological replicates are shown as black and white circles.

### OmpF is beneficial but not essential for CdiA^F11^ mediated growth inhibition

Another factor that could affect effector delivery, and which could be different on LB than on M9Glu solid media, is the receptor abundance. The expression of the OmpC/OmpF receptor proteins are inversely regulated by osmolarity through a single regulator OmpR, which slightly represses the expression of OmpC and strongly increases the expression of OmpF in response to a decrease in osmolarity (Forst, Delgado, Ramakrishnan, & Inouye, 1988). As LB has a higher NaCl concentration than M9Glu, less OmpF should be expressed on this media, resulting in less OmpC/OmpF on the cell surface. We therefore investigated if OmpC/OmpF expression differs between LB and M9Glu media by western blot. MG1655 cells expressed very similar levels of OmpC during growth in all tested conditions (Fig. 4A), whereas OmpF expression increased during growth in low salt media (M9Glu or LB with low concentrations of NaCl) (Fig. 4B).

**Figure 4.**
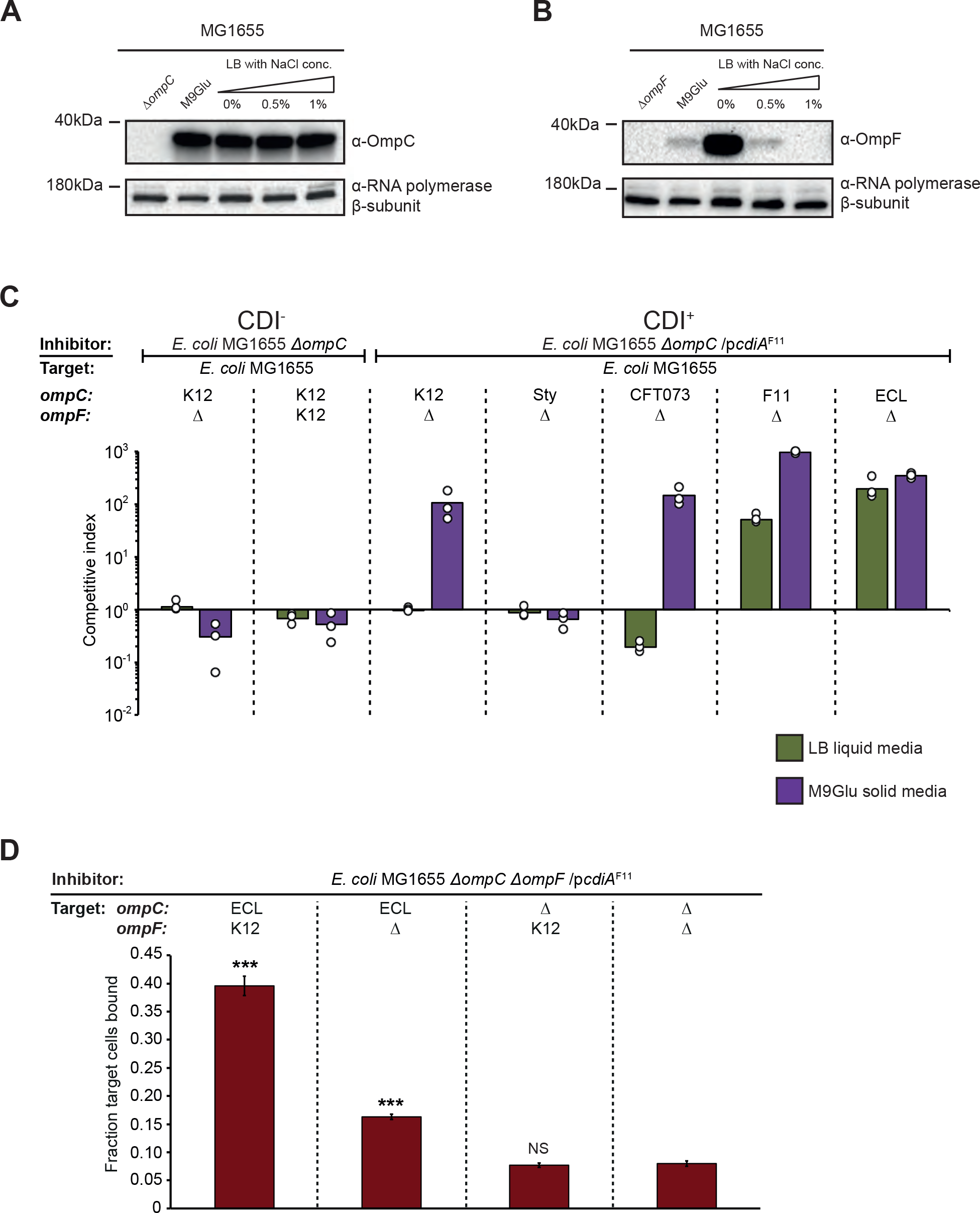
OmpF is not required for class II CdiA mediated growth inhibition. **A)** OmpC expression from the MG1655 chromosome measured by Western blot, probed with an anti-OmpC antibody. RNA polymerase *β*-subunit was used as loading control. **B)** OmpF expression from the MG1655 chromosome measured by Western blot, probed with an anti-OmpF antibody. RNA polymerase *β*-subunit was used as loading control. **C)** Average competitive index of cells expressing CdiA^F11^ after co-culturing with Δ*ompF* MG1655 cells expressing different OmpC’s from the native MG1655 *ompC* locus (n=3 biological replicates). Cells were co-cultured for 5h in liquid LB media (green bars) or 24h on solid M9Glu media (purple bars). Individual data points of the biological replicates are shown as black and white circles. **D)** Average fraction of MG1655 target cells with or without *ompC*ECL and *ompF*K12 bound to CdiA^F11^+ inhibitor cells. Error-bars are SEM. Statistical significance was determined using a two-tailed, unpaired students t-test where * = P<0.05, ** = P<0.005 and ***= P<0.0005.

From these results, it appeared as if CdiA^F11^ effector delivery could occur even when OmpF levels were very low (undetectable by western blot) (Fig 4B; LB 1% salt is the condition that the competitions in Fig. 1A were performed in), making us question whether OmpF is required for CdiA^F11^ mediated growth inhibition as previously demonstrated (Beck et al., 2016). To test if OmpF was required for class II CdiA effector delivery we competed cells expressing CdiA^F11^ with cells expressing K12, *Sty*, CF073, F11 or ECL OmpC, in the absence of OmpF. Interestingly, *ΔompF* cells expressing either OmpC^F11^ or OmpC^ECL^ were outcompeted by CdiA^F11^ expressing cells when grown in liquid LB (Fig. 4C), suggesting that OmpF is not required for CdiA^F11^ mediated growth inhibition. In addition, *ΔompF* cells expressing any of the investigated OmpC variants, with the exception of OmpC^Sty^, were inhibited on M9Glu solid media (Fig. 4C), suggesting that OmpF is not essential for class II CdiA effector delivery, but that it is beneficial when the CdiA-OmpC binding interaction is weak. To investigate if OmpF mediated an increase in relative cell-cell binding in liquid LB media we measured inhibitor-target cell interactions between inhibitor cells expressing CdiA^F11^ and target cells expressing OmpC^ECL^ in the presence or absence of OmpF using a modified cell-cell binding assay. To capture even a weak CdiA-receptor interaction, inhibitor and target cells were cross-linked by formaldehyde after 40 min of co-cultivation. The relative inhibitor-target cell interactions were measured by flow cytometry as previously described (Fig. 2A-B). A significantly larger fraction (40%) of target cells expressing OmpC^ECL^ and OmpFK12 were bound to inhibitor cells expressing CdiA^F11^, as compared to target cells only expressing OmpC^ECL^ (15%) (Fig. 4D). These results further confirm that although OmpF is not essential for cell-cell binding or toxin delivery, it stabilizes the CdiA-receptor interaction.

### Class II CdiA effector translocation has not evolved to favor intra-species interactions

Our results suggest that CdiA^F11^ has a higher binding affinity and preference for the OmpC^F11^ and OmpC^ECL^ receptors over others. We therefore wanted to investigate if the minor differences observed between the RBD of CdiA^F11^, CdiA^CFT073^ and CdiA^Sty^ (Fig. S1), create an intra-species preference of receptor binding, i.e. does CdiA^CFT073^ favor a binding to OmpC^CFT073^, and CdiA^Sty^ a binding to OmpC^Sty^ ? To this end we created chimeric CdiA proteins where the CdiA receptor-binding domain of CdiA^F11^ was changed to that of *S. typhi* (CdiA^Sty^) or *E. coli* CFT073 (CdiA^CFT073^). Cells expressing CdiA^Sty^ did not outcompete cells expressing low levels of OmpC^Sty^ better than cells expressing other OmpC variants in liquid LB media (Fig. 5A). Similarly, cells expressing CdiA^CFT073^ did not outcompete cells expressing low levels of OmpC^CFT073^ better than cells with any other OmpC variant (Fig. 5B), instead both CdiA^Sty^ and CdiA^CFT073^ outcompeted cells expressing OmpC^ECL^ the most (Fig. 5A-B). Furthermore, all target cells expressing any of the different OmpC variants were inhibited on M9Glu solid media when co-cultivated with inhibitor cells expressing either CdiA^Sty^ or CdiA^CFT073^ (Fig. S5). Thus, the general trend of growth inhibition for both CdiA^Sty^ and CdiA^CFT073^ is identical to that of CdiA^F11^. Taken together this suggests that the minor differences found in the receptor-binding domain of different class II CdiA proteins do not result in preferentially targeting of intra-species receptors.

**Figure 5.**
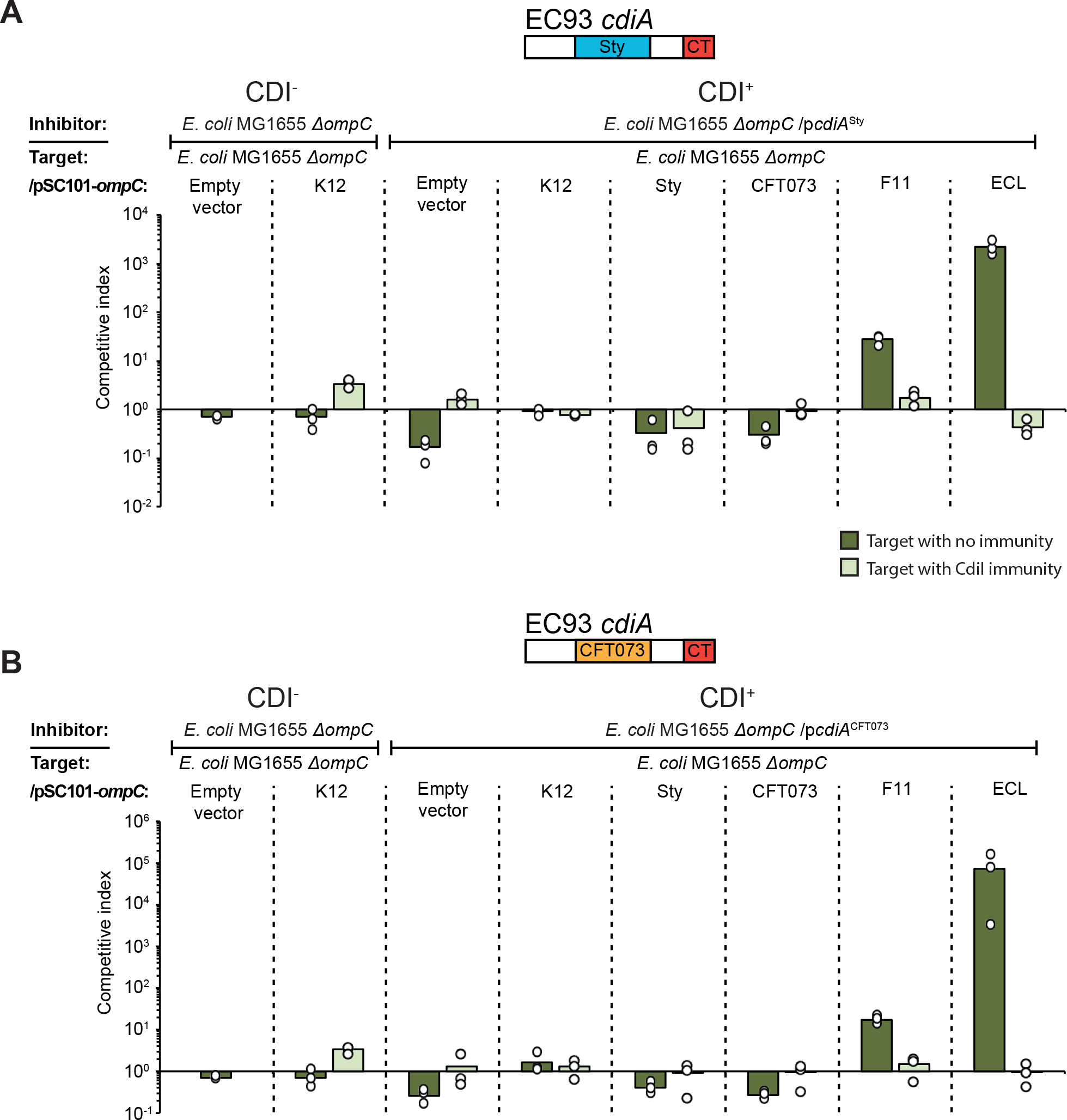
Other Class II CdiA RBD is able to deliver effectors to cells expressing the OmpC receptor from other strains and species. Average competitive index of cells expressing **A)** CdiA^Sty^ or **B)** CdiA^Nissle^ after co-culturing with MG1655 cells expressing different OmpC’s from a constitutive PJ23101 promoter on a low-copy (pSC101) plasmid (n=3 biological replicates). Cells were co-cultured for 5h in liquid LB media. Individual data points of the biological replicates are shown as black and white circles.

### Class II CdiA mediated CDI works cross-species

Our results showed that CdiA mediated growth inhibition of *E. coli* target cells expressing OmpC’s from other species is possible. This does not necessarily infer that *E. coli* cells expressing CdiA^F11^ can actually inhibit the growth of other species. To test if actual strains of *E. cloacae* (ATCC 13047)*, S. typhimurium* LT2, UPEC 536 and *E. coli* CFT073 were inhibited by CdiA^F11^ mediated toxin delivery, we competed *E. coli* cells expressing CdiA^F11^ with these bacterial species. In addition, we extended the set of bacteria by including other species where OmpC homologs could be identified bioinformatically (*Enterobacter aerogenes* and *Klebsiella pneumoniae*) (Fig. S6). To control for other factors e.g. differences in growth or delivery of other toxins that could affect the competition, we provided these bacterial species with CdiI immunity proteins expressed from a medium-copy plasmid. In liquid LB media, *E. coli* MG1655 cells expressing CdiA^F11^ outcompeted wild-type *E. cloacae* and *E. aerogenes* cells with 2-logs, whereas all other tested wild-type bacterial strains were not outcompeted (Fig. 6A, dark green bars). On M9Glu solid media, *E. coli* MG1655 cells expressing CdiA^F11^ outcompeted *E. coli* CFT073, UPEC 536 and *E. aerogenes* wild-type cells with 2- to 3-logs (Fig. 6B, dark purple bars), whereas the others were not outcompeted (Fig. 6B). As expected, cells complemented with *cdiI* were not outcompeted on either media (Fig. 6A-B, light colored bars). Interestingly, *E. cloacae* cells were inhibited in liquid LB but not on solid M9Glu media. Previous results suggest that *E. cloacae* harbors a T6SS active against *E. coli* on solid media (Beck et al., 2014). Thus, we hypothesized that our CdiA^F11^ expressing *E. coli* could be inhibited back by *E. cloacae* on solid M9Glu media. To test if this was the case we competed *E. coli* MG1655 cells expressing CdiA^F11^ against a Δ*vasK* mutant (unable to form the T6SS) of *E. cloacae. E. coli* MG1655 cells expressing CdiA^F11^ outcompeted *E. cloacae* cells lacking *vasK* with 2-logs on both liquid and solid media, suggesting that the ability to compete back could indeed explain the differential inhibition on solid and liquid media. Similarly, *S. typhimurium* cells complemented with *cdiI* outcompeted MG1655 inhibitor cells by 1-log indicating that *S. typhimurium* cells were indeed inhibited by MG1655 inhibitor cells but that differences in other fitness factors were hiding this fact (Fig. 6B). To normalize against any unknown fitness factors we transformed wild-type *S. typhimurium* cells with our CdiA^F11^ expressing plasmid and competed these cells against wild-type *S. typhimurium* (Fig. 6C). *S. typhimurium* cells expressing CdiA^F11^ outcompeted wild-type *S. typhimurium* by almost 2-log on M9Glu solid media but not in liquid LB (Fig. 6C). This clearly showed that *S. typhimurium* can both inhibit and be inhibited by a class II CDI system (CdiA^F11^).

**Figure 6.**
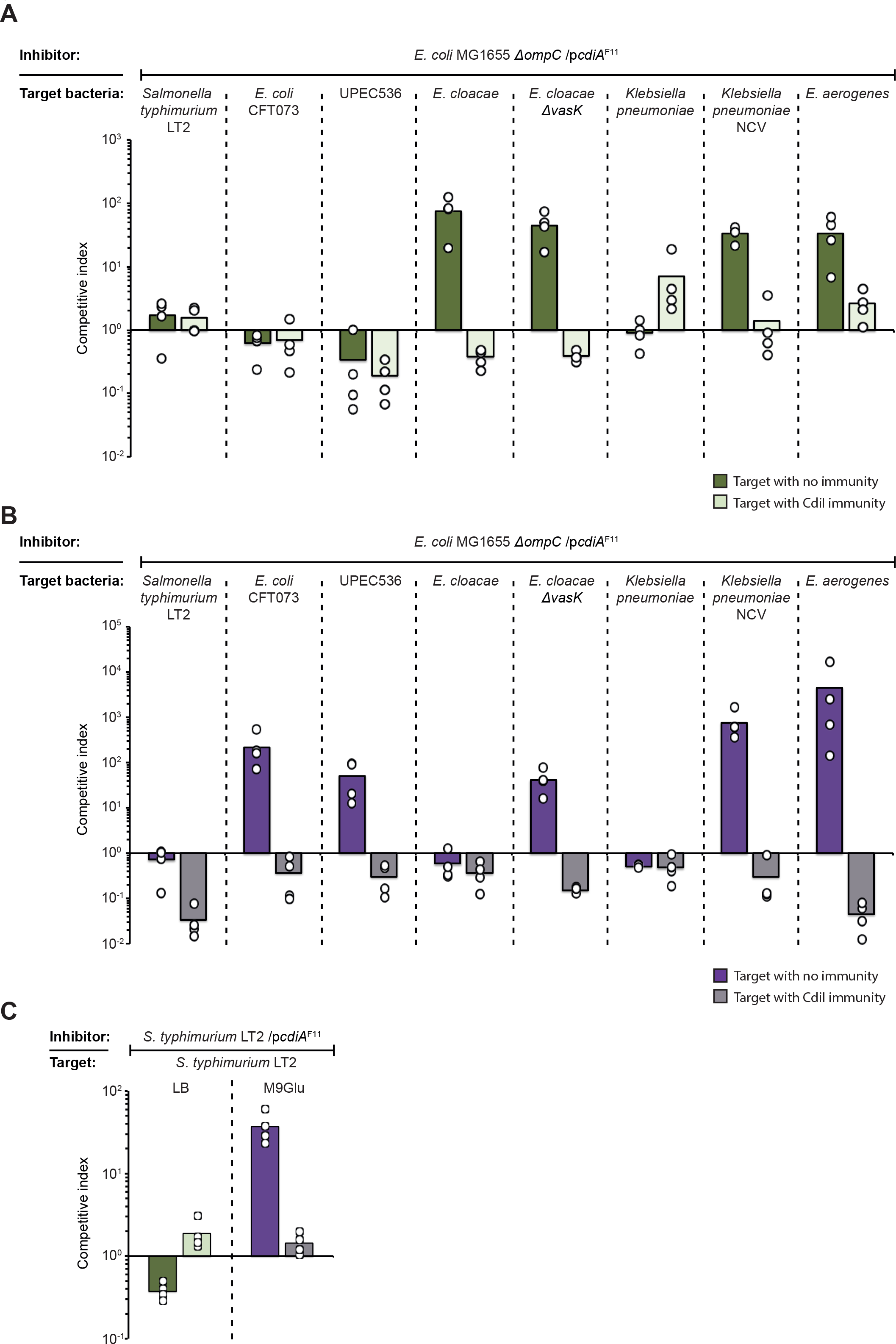
Class II CdiA toxin delivery works cross-species. **A & B)** Average competitive index of cells expressing CdiA^F11^ after co-culturing with *Salmonella typhimurium E. coli* CFT073*, E. coli* UPEC 536, *Enterobacter cloacae*, *Klebsiella pneumoniae* or *Enterobacter aerogenes* with (light green bars) or without (dark green bars) CdiI expressed from plasmid. Co-culturing was performed for 5h in liquid LB media (**A**) or 24h on solid M9Glu media (**B**) (n=4 biological replicates). **C)** CdiA^F11^ expressing *S. typhimurium* LT2 competed against *S. typhimurium* LT2 with (light bars) or without (dark bars) CdiI immunity on liquid LB (green) or solid M9Glu (purple) media (n=4 biological replicates). Individual data points of the biological replicates are shown as black and white circles.

Furthermore, no growth inhibition could be observed against *K. pneumoniae* when competed against *E. coli* MG1655 cells expressing CdiA^F11^, whereas *E. aerogenes* was outcompeted on both media (Fig. 6A-B). From the colony morphology it was obvious that the *K. pneumoniae* strain NC105 was forming a lot of capsule (not shown). We therefore examined if *E. coli* cells expressing CdiA^F11^ were able to outcompete a non-capsulated mutant of *K. pneumonia. E. coli* MG1655 cells expressing CdiA^F11^ outcompeted capsule deficient *K. pneumoniae* by 2-log in liquid and on solid media showing that capsule is an obstacle for CdiA^F11^ mediated toxin delivery (Fig. 6A-B).

To verify that effector delivery can occur by OmpC homologs from *K. pneumoniae* and *E. aerogenes*, we cloned the OmpC homologs into the same pSC101 plasmid used to express all other OmpC variants in this study. *E. coli* MG1655 cells expressing CdiA^F11^ outcompeted *E. coli* cells expressing OmpC from *K. pneumoniae* and *E. aerogenes* with 2-logs (Fig. S7), showing that CdiA^F11^ mediated effector delivery was possible with these OmpC homologs. Taken together, these results suggest that although other factors limit CDI, class II CdiA^F11^ mediated growth inhibition can occur cross-species.

### CdiA from *Enterobacter cloacae* is an *E. coli* class II CdiA analog

This is not the first study showing cross-species inhibition by CdiA. A previous study looking at CdiA mediated toxin delivery in *E. cloacae* showed that when the CdiA protein of *E. cloacae* (CdiA^ECL^) was artificially over-expressed from an arabinose inducible promoter, *E. cloacae* strains unable to utilize its T6SS (*ΔvasK*) could still inhibit the growth of *E. coli*, suggesting that CDI could work cross-species (Beck et al., 2014). As our results suggest that class II CdiA RBD’s from *E. coli* are promiscuous and capable of delivering CdiA effectors to cells expressing OmpC proteins from other species, we were interested to investigate if the *E. cloacae* CdiA RBD was also promiscuous. The CdiA RBD of CdiA^ECL^ has no sequence homology to the RBD of CdiA^F11^ (Fig. S8), so we first looked for the receptor of the *E. cloacae* CdiA protein by creating a chimeric CdiA protein where the CdiA receptor-binding domain of CdiA^F11^ was changed to that of *E. cloacae* CdiA (CdiA^ECL^). Next, we created a mariner transposon pool in a MG1655 strain expressing *acrB* (known permissive factor of the EC93 CdiA ionophore toxin used in all our constructs) from a multi-copy plasmid and enriched for resistant mutants against CdiA^ECL^ by repeatedly competing the transposon pool with MG1655 cells expressing CdiA^ECL^. After three rounds of enrichment, we isolated resistant mutants and used semi-random arbitrary PCR to identify the insertion sites of the transposons providing resistance towards CdiA^ECL^ toxin delivery. Interestingly, we identified one transposon located 3nt upstream of the *ompF* ORF, and one 630bp in the *ompF* ORF (Fig. 7A), suggesting that OmpF alone or OmpC/OmpF heterotrimers function as the receptor(s) for CdiA^ECL^. To verify that CdiA^ECL^ was also using the OmpC/OmpF heterotrimers as a receptor we competed *ΔompC* or *ΔompF* mutants of *E. coli* MG1655 against inhibitor cells expressing CdiA^ECL^. Cells expressing CdiA^ECL^ outcompeted MG1655 target cells with 1-log on M9Glu solid media (Fig. 7B), but could not outcompete *E. coli* cells lacking either *ompC* or *ompF* (Fig. 7B), confirming that OmpC/OmpF heterotrimers function as the receptor for CdiA^ECL^.

**Figure 7.**
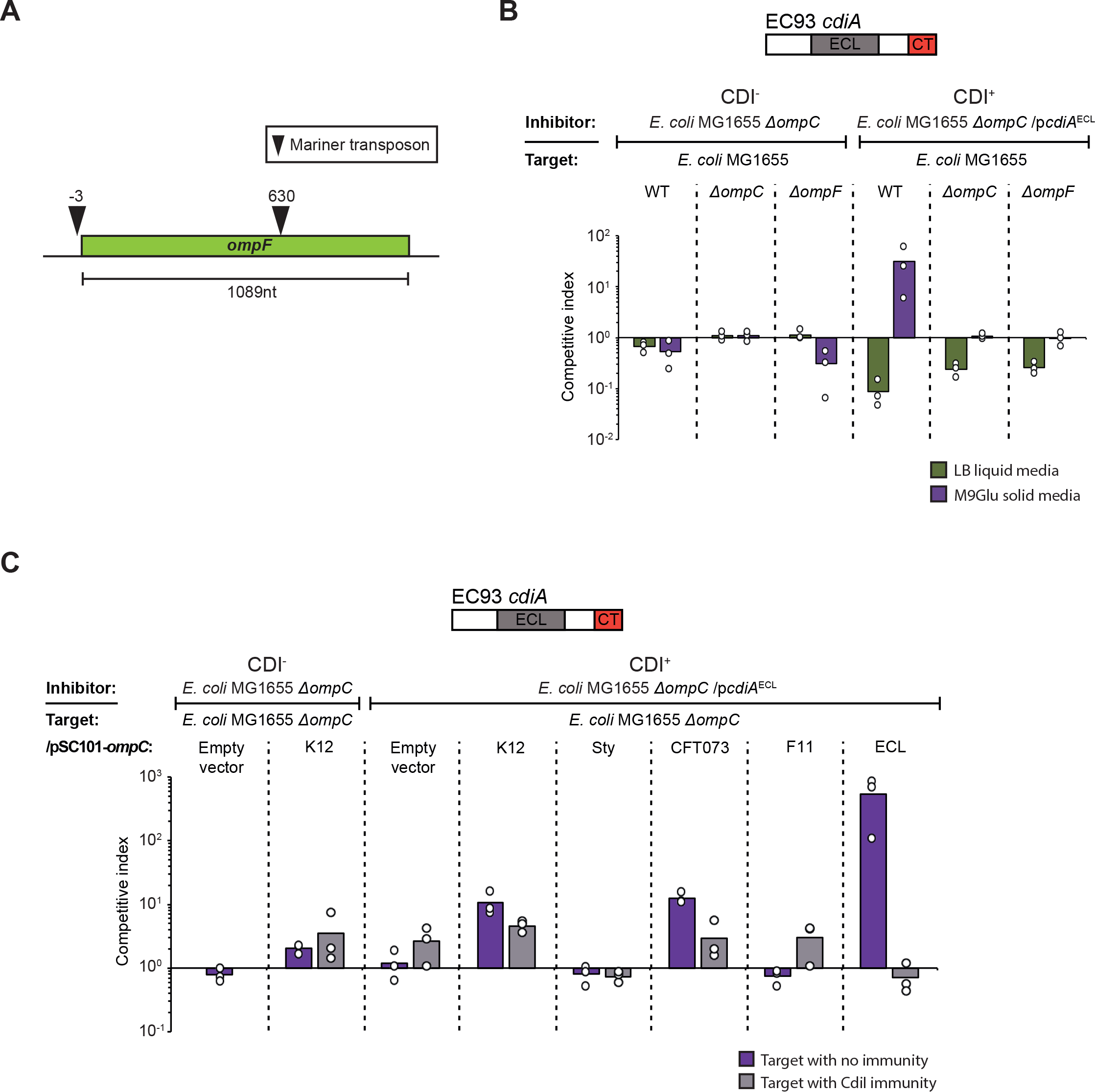
The CdiA protein of *E. cloacae* is a Class II CdiA analog. **A)** Illustration of the identified mariner transposon insertion sites of CdiA^ECL^ resistant target cells. **B & C)** Average competitive index of cells expressing CdiA^ECL^ after co-culturing with MG1655 cells lacking OmpC/F (**B**) or expressing different OmpC’s from a constitutive PJ23101 promoter on a low-copy (pSC101) plasmid (**C**). Co-culturing was for 5h in liquid LB media (green bars) or for 24h on solid M9Glu (purple bars) media (n=3 biological replicates). Individual data points of the biological replicates are shown as black and white circles.

As CdiA^ECL^ required the same receptor complex as CdiA^F11^, while sharing little to no sequence homology with CdiA^F11^, we wanted to know if CdiA^ECL^ was an equally promiscuous class II CdiA analog and thus also able to inhibit cells expressing other OmpC receptors. To test this we competed cells expressing CdiA^ECL^ against MG1655 cells with *ompC* from *E. cloacae*, K12, Sty, CFT073 or UPEC F11. Inhibitor cells expressing CdiA^ECL^ outcompeted cells expressing low levels of OmpC^ECL^ by >2-logs, and OmpC^K12^ or OmpC^CFT073^ by 1-log on solid M9Glu media (Fig. 7C). Cells expressing low levels of OmpC^F11^ or OmpC^*Sty*^ were not outcompeted by cells expressing CdiA^ECL^ (Fig. 7C) and none of the targets were outcompeted in liquid media (Fig. S9). Taken together, these results show that CdiA^ECL^ is a class II CdiA analog that is also able to deliver effectors cross-species.

## Discussion

Here we show that class II CdiA proteins from *E. coli* allow for efficient effector delivery to a range of *Enterobacteriaceae* spp. including *E. coli*, *S. typhimurium*, *E. cloacae*, *E. aerogenes* and *K. pneumoniae*. In contrast to previous findings (Beck et al., 2016), our results show that OmpC receptor-binding by the class II CdiA molecules and their *E. cloacae* CdiA analog is very promiscuous, allowing all four tested CdiA chimeras to deliver effectors cross-species without any intra-species or strain preference. These findings are supported by previous studies in other species. For example, over-expression of a naturally occurring CdiBAI system in *E. cloacae* allows inhibition of *E. coli* (Beck et al., 2014) and *Burkholderia pseudomallei* CdiA delivers effectors to *Burkholdera thailandensis* (Koskiniemi et al., 2015). Broad-range toxin delivery to other species that share the same growth niche could be beneficial for antagonistic interactions as well as for kin recognition. The ability to recognize siblings from other bacteria is important in any community that exchanges resources. For kin recognition, two types of recognition are important: i) to identify the own kin from others in order to bind and exchange outer-membrane vesicles only with kin cells, as has been shown to occur by the TraA and IdsD/E systems in *Myxococcus xanthus* and *Proteus mirabilis* respectively (Gibbs, Urbanowski, & Greenberg, 2008; Pathak, Wei, Dey, & Wall, 2013). And ii) inhibition of non-kin bacteria that would otherwise utilize shared resources, as has been shown to occur by T6SS mediated toxin delivery in *P. mirabilis* or *M. xanthus* (Vassallo et al., 2017; Wenren, Sullivan, Cardarelli, Septer, & Gibbs, 2013). For the former, very specific protein-protein interactions are favorable and only a few amino acid differences can block binding completely (P. Cao & Wall, 2017). Whereas for the latter, a broader specificity is desirable as it allows inhibition of also more distant unrelated species that could utilize the same resources. CdiA proteins are versatile in that they both allow tight cell-cell binding and toxin delivery (Aoki et al., 2010; Aoki et al., 2005; Ruhe et al., 2015). But how CDI contributes to kin recognition, i.e. if it is by identifying self from others or by inhibiting the growth of other bacteria, is not known. Our results suggest that class II CdiA proteins function through the latter, where promiscuous effector delivery allows for a broader range of non-kin recognition and subsequent growth inhibition.

The remaining question is why class II CdiA proteins are different from class I CdiA proteins, where no promiscuous effector delivery can be observed (Ruhe, Wallace, et al., 2013)? Class I and II CdiA molecules bind and recognize different receptor proteins, so a possible explanation could be the abundance of the two receptor-proteins on the surface of bacterial cells. OmpC is the most abundant protein on the bacterial cell surface, expressed at approximately 100,000 molecules/cell (Schuman, 2006). As a consequence, OmpC proteins are under strong selection to undergo immunogenic variation to avoid targeting by the host immune system (Liu et al., 2012; Singh, Williams, Klebba, Macchia, & Miller, 2000; Stenkova, Bystritskaya, Guzev, Rakin, & Isaeva, 2016). In addition, OmpC is the receptor for numerous phages, increasing the selection pressure for constant change to avoid phage infections (Bertozzi Silva, Storms, & Sauvageau, 2016). This selection for change is easily observable in the extracellular loops of OmpC proteins that vary extensively between different *E. coli* strains (Fig. S2 and S6). Such variations can only be observed between species for BamA, which although also functions as a phage receptor, is expressed more than one order of magnitude less (approximately 4000 molecules/cell (Li, Burkhardt, Gross, & Weissman, 2014). Thus, one possibility is that the species-promiscuity of the class II receptor-binding domains is a consequence for retaining self-or intra-species delivery, which might be important for the ability to recognize the most important competitors of the own niche, i.e. those very similar but not kin. On the other hand, self-delivery of CdiA effectors (delivery of toxins to cells with cognate immunity) was recently shown to be important for contact-dependent signaling and in increasing stress tolerance (Garcia, Perault, Marlatt, & Cotter, 2016; Ghosh et al., 2018), suggesting that CDI could be used for both cell-cell communications as well as for antagonistic interactions and kin recognition. Thus, retaining self-or intra-species delivery could be important for functional cell-cell communication as well as for kin-cell recognition.

A final question that arises is if CDI is important in shaping bacterial communities and if so whether it plays a role during initial colonization of a niche and/or in protecting an established micro-colony. In an aqueous environment with flow (like the host gut), contact-mediated effector delivery and ability for tight adherence to the targeted cell should be favored over secretion of diffusible toxins to the extracellular milieu, as has been suggested for the glycine zipper like protein toxins from the freshwater bacteria *Caulobacter cresentus* (Garcia-Bayona, Guo, & Laub, 2017). Thus, it is possible that a higher binding affinity for certain receptors allows CDI positive cells to adhere and deliver effectors to their main competitors of this niche during initial contact in the host gut and that this is more important for some bacteria than others. For example, we find that CdiA proteins from UPEC 536 and *E. cloacae* have a higher binding affinity to the own receptor over others, but that this is not the case for the CFT073 and *S. typhi* homologs. Once the micro-colony has been established however, contacts with new invading species or strains will mainly occur at the edges of the growing colony, where toxin delivery occur as efficiently cross-species as within species. In this context, class II CDI seems to function as a kin recognition system, where antagonistic interactions with a broad range of other species frequently found in the niche, could potentially allow bacteria with CDI to protect the borders of the growing micro colony against foreign attacks and to restrict the use of shared resources to the own kin. Taken together, our results suggest that class II CdiA proteins are versatile molecular machineries that allow for different behavior depending on the environmental context.

## Methods

### Strains and growth conditions

The bacterial strains used in this study are listed in Table S1. Strains were grown at 37°C and 200 rpm shaking in Luria-Bertani broth, LB, (10 g/l Tryptone, 5 g/l Yeast extract and 10 g/l NaCl) unless stated otherwise. M9 minimal medium (1x M9 salts, 2 mM MgSO_4_, 0.1 mM CaCl_2_) was supplemented with 0.4% glucose, 0.2% casamino acids and 50 mM FeCl_3_. Media were supplemented with antibiotics when applicable as follows: ampicillin (AMP) 100 mg/l, chloramphenicol (CAM) 12.5 mg/l, kanamycin (KAN) 50 mg/l, streptomycin (STREP) 100 mg/l and Spectinomycin (SPEC) 50 mg/l.

### Construction of plasmids and chromosomal constructs

Gene deletions were retrieved from the Keio collection (Baba et al., 2006) and moved between strains by P1 transduction. All constructs were verified by PCR and sequencing. For detailed information of the different constructs see the supplementary material.

### Competition assay

The cells were grown overnight in LB. Inhibitor and target cells were mixed at a ratio of 10:1 and either diluted 1:100 in LB for liquid media competitions or 20 μl were spotted on solid M9Glu minimal media. For liquid media competitions, the cells were co-cultured for 5 h at 37°C with 200 rpm shaking and plated onto LB solid media containing appropriate antibiotics to enumerate inhibitor and target cells as the number of colony-forming units per milliliter (CFU/ml). For competitions on solid media, the cells were co-cultured for 24 h at 37°C before suspended in 1xPBS, followed by enumeration of inhibitor and target cells as above. Competitive indexes were calculated as the ratio of inhibitor to target cells at the end of the co-culture (5 h or 24 h) divided by the ratio at the beginning of the co-culture. The competitive indexes for three independent experiments are reported ± standard error of the mean.

### Membrane-protein enrichment and SDS-PAGE to analyze OmpC

MG1655 cells expressing different variants of OmpC from the chromosome or from a low-copy plasmid (table S1) were diluted 1/1000 from an over-night (ON) culture and grown to stationary phase (OD600 = 2.0). 1 ml of each bacterial culture were pelleted at 21000 x g for 10 min and re-suspended in 2ml of a mild hypotonic lysis buffer (50 mM Tris pH6.8, 1% Triton X-100, 1 mg/ml lysozyme, 10 mM EDTA, 1 tablet/50ml SIGMAFAST Protease Inhibitor Cocktail (Sigma-Aldrich, Germany)) and incubated at room-temperature for 60min. Cells were then subjected to 6 cycles of freeze-thawing in a ethanol dry-ice bath before being pelleted at 21000 x g for 60 min. Supernatants were discarded and pellets was re-suspended in 2 ml wash buffer (50 mM Tris pH6.8, 2 mM MgSO_4_, 10 U/ml Benzonase (Sigma-Aldrich, Germany)) and incubated an additional 20min on ice to degrade genomic DNA. Cells were then pelleted again at 21000 x g for 60 min and the pellet was re-suspended in 100μl of a 1x membrane-protein sample buffer (50mM Tris pH6,8, 1% SDS (w/v), 1% Triton X-100, 10% Glycerol, 0.2% bromophenol blue) and boiled for 5min at 95°C. 150mM DTT was then added to each sample followed by the pelleting of membrane debris and undigested genomic DNA at 21000 x g for 5 min before the supernatant were separated on a Mini-PROTEAN TGX protein gel (Biorad, USA). Total protein was detected in gel by SYPRO Ruby Protein Gel Stain (Thermo Scientific, USA) and visualized by UV-table.

### Western blot to analyze OmpC and OmpF

500 μl of an ON-culture of MG1655 cells were harvested by centrifugation for 5 min at 21000 x g. The supernatant was discarded, and cells were re-suspended in 250 μl of 1x membrane-protein sample buffer (50mM Tris pH6,8, 1% SDS (w/v), 1% Triton X-100, 10% Glycerol, 0.2% bromophenol blue) and boiled for 5min at 95°C. 150mM DTT was then added to each sample followed by the pelleting of membrane debris and undigested genomic DNA at 21000 x g for 5 min before the supernatant were separated on a Mini-PROTEAN TGX gel (Biorad, USA). PageRuler Prestained Protein Ladder (Thermo Scientific, USA) was used as size marker. Proteins were then transferred to a Trans-Blot Turbo Mini 0.2 μm PVDF membrane (BioRad, USA) using the Trans-Blot Turbo system (Biorad, USA). OmpC proteins were detected using anti-OmpC antibody (orb308739, Biorbyt, United Kingdom) and OmpF proteins were detected using anti-OmpF antibody (orb308741, Biorbyt, United Kingdom). Equal loading was confirmed by probing for RNA polymerase β-subunit using an anti-RNAP β-subunit antibody (ab191598, Abcam, United Kingdom). Secondary antibody towards anti-OmpC, anti-OmpF and anti-RNAP antibodies was anti-rabbit IgG coupled to HRP (A1949, Sigma-Aldrich, Germany). Bands were visualized with Clarity Western ECL Substrate (Biorad, USA) and a ChemiDoc MP system (Biorad, USA).

### Flow cytometry cell-cell binding assay without cross-linking

MG1655 *ΔompC* inhibitor cells with a galK::sYFP2-catR casett integrated on the chromosome and target cells with a galK::dTomato-catR casett, also integrated on the chromosome (both fluorophores expressed by the synthetic iGEM promoter pJ23101) (table S1) were diluted 1/100 into LB from an ON-culture and grown to stationary phase (OD600 = 2.0). Inhibitor and target cells were then mixed at a 5:1 ratio and incubated for 40 min with aeration at 37°C. Samples were then diluted 1/1000 in 1xPBS followed by a gently mixing before samples were analyzed by a MACSQuant VYB flow cytometer using filters B1 (525/50nm) and Y2 (615/20nm) (Miltenyi Biotec). Flow rate was adjusted to allow for 1500 events/sec and at least 100 000 events were collected. Fraction of dTomato-sYFP2 double events were analyzed in relation to total number of dTomato events (fraction target cells bound to a inhibitor) with FlowJo Software (FlowJo, LLC, USA).

### Flow cytometry cell-cell binding assay with cross-linking

MG1655 *ΔompC* inhibitor cells with a galK:: dTomato-catR casett integrated on the chromosome and target cells transformed with a plasmid expressing sYFP2 (both fluorophores expressed by the synthetic iGEM promoter pJ23101) (table S1) were diluted 1/100 into LB from an ON-culture and grown to stationary phase (OD600 = 2.0). Inhibitor and target cells were then mixed at a 5:1 ratio and incubated for 40 min without agitation at 37°C after which 4% (final conc.) formaldehyde was added and samples were incubate in room-temp. for an additional 20 min. Samples were then diluted 1/1000 in 1xPBS and vortexed heavily for 10 sec before being analyzed by a MACSQuant VYB flow cytometer, same as above. Flow rate was adjusted to allow for 500 events/sec and at least 25000 events were collected. Fraction of sYFP2-dTomato double events (fraction target cells bound to a inhibitor) were analyzed in relation to total number of sYFP2 events with FlowJo Software (FlowJo, LLC, USA).

## Acknowledgments

This study was supported by grants from the Swedish Foundation of Strategic Research, the Swedish research council, Uppsala antibiotic center and the Åke Wiberg and Wenner-Gren foundations (to S.K).

## Data availability

All data included in the manuscript will be made publicly available if accepted for publication and will be made available for reviewers upon request.

## Supplementary figure legends

**Figure S1.**
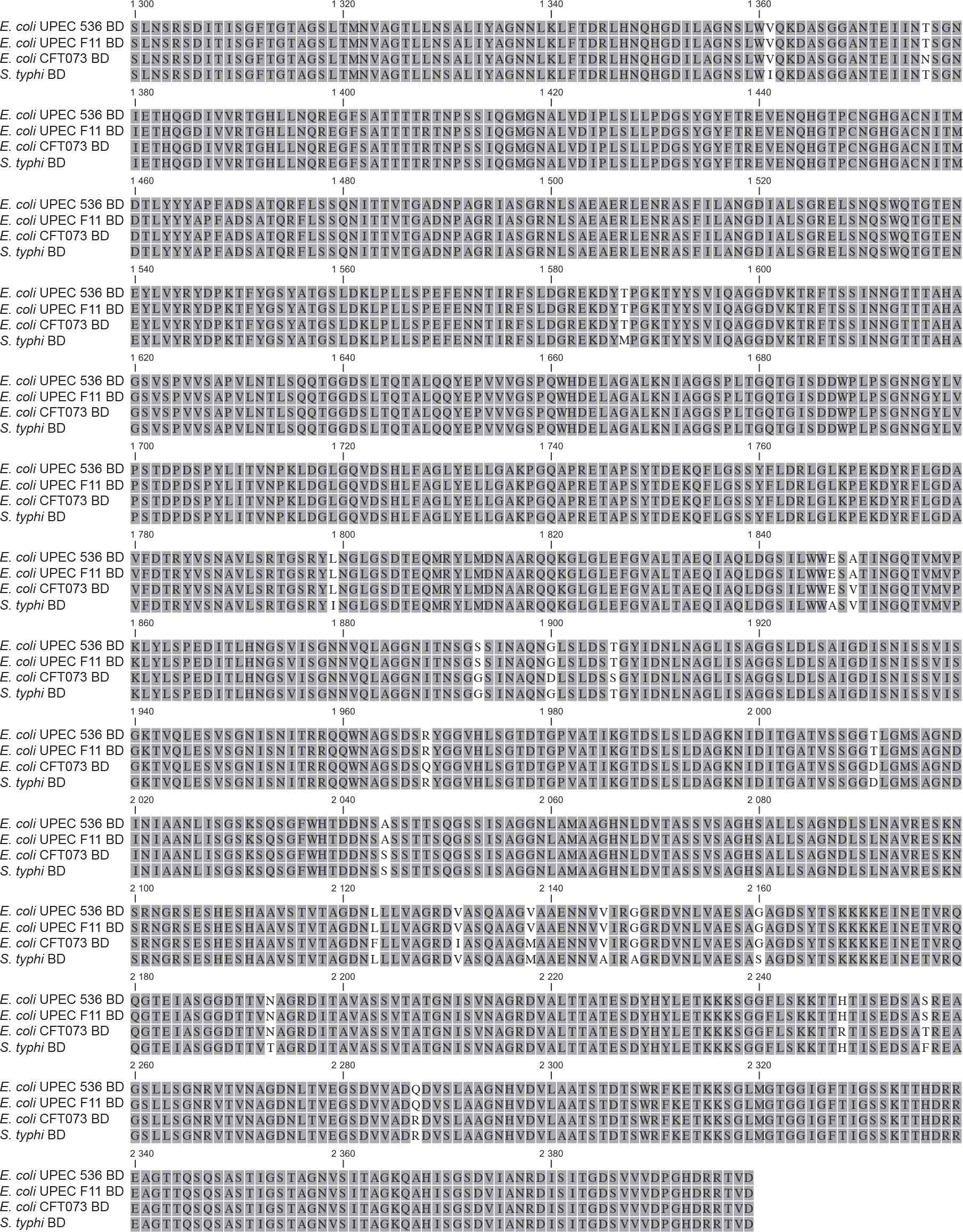
Alignment of CdiA receptor binding domains from *Escherichia coli* strains UPEC 536, UPEC F11, CFT073 and *Salmonella typhi*. Homologous residues are shown in grey and non-homologous residues are white.

**Figure S2.**
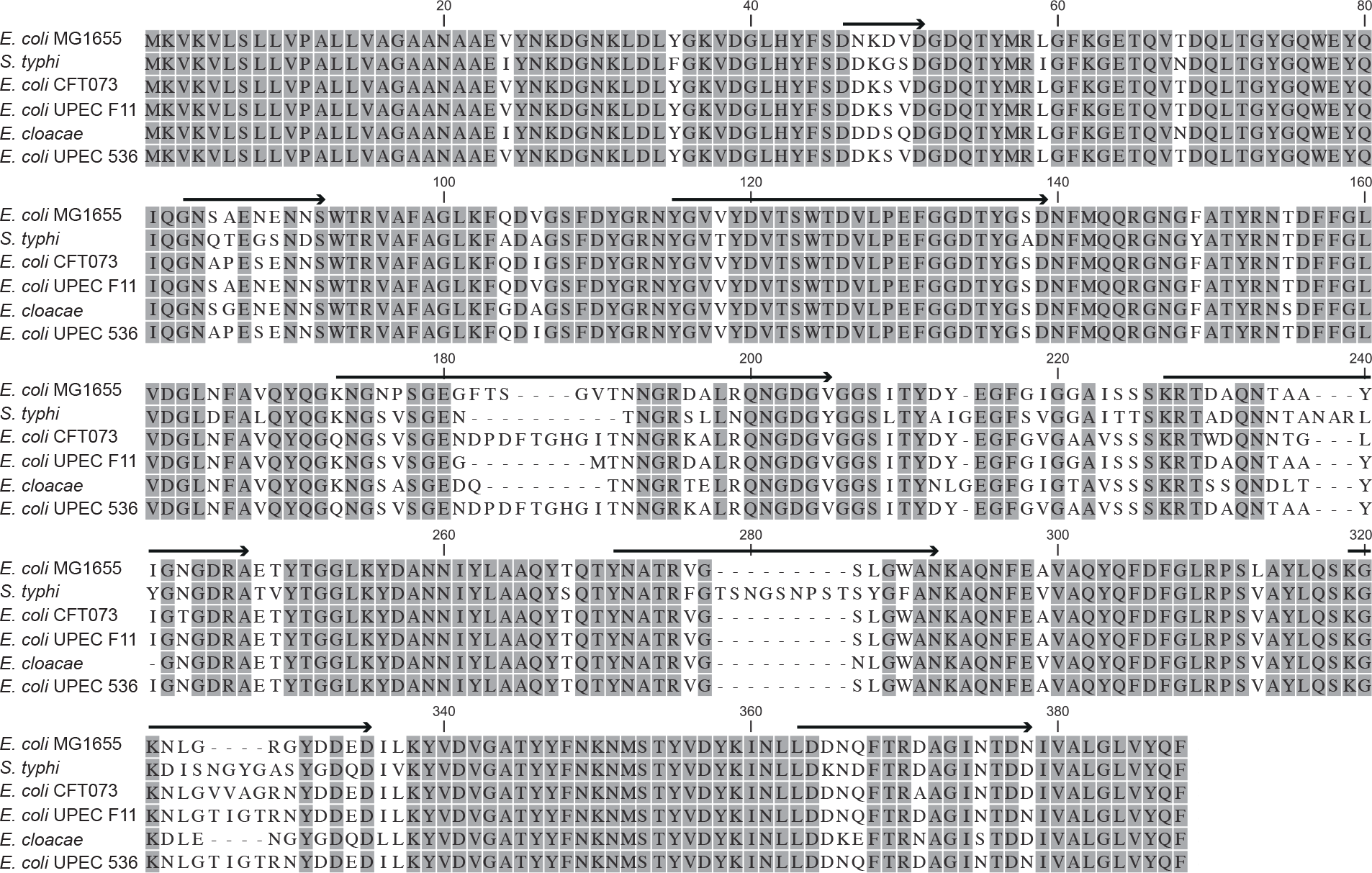
Alignment of OmpC proteins from *Escherichia coli* strains K12 MG1655, UPEC 536, UPEC F11, CFT073 and *Enterobacter cloacae and Salmonella typhi*. Homologous residues are shown in grey and non-homologous residues are white. The location of the 8 extracellular loops in OmpC are marked with a black arrow above the sequence.

**Figure S3.**
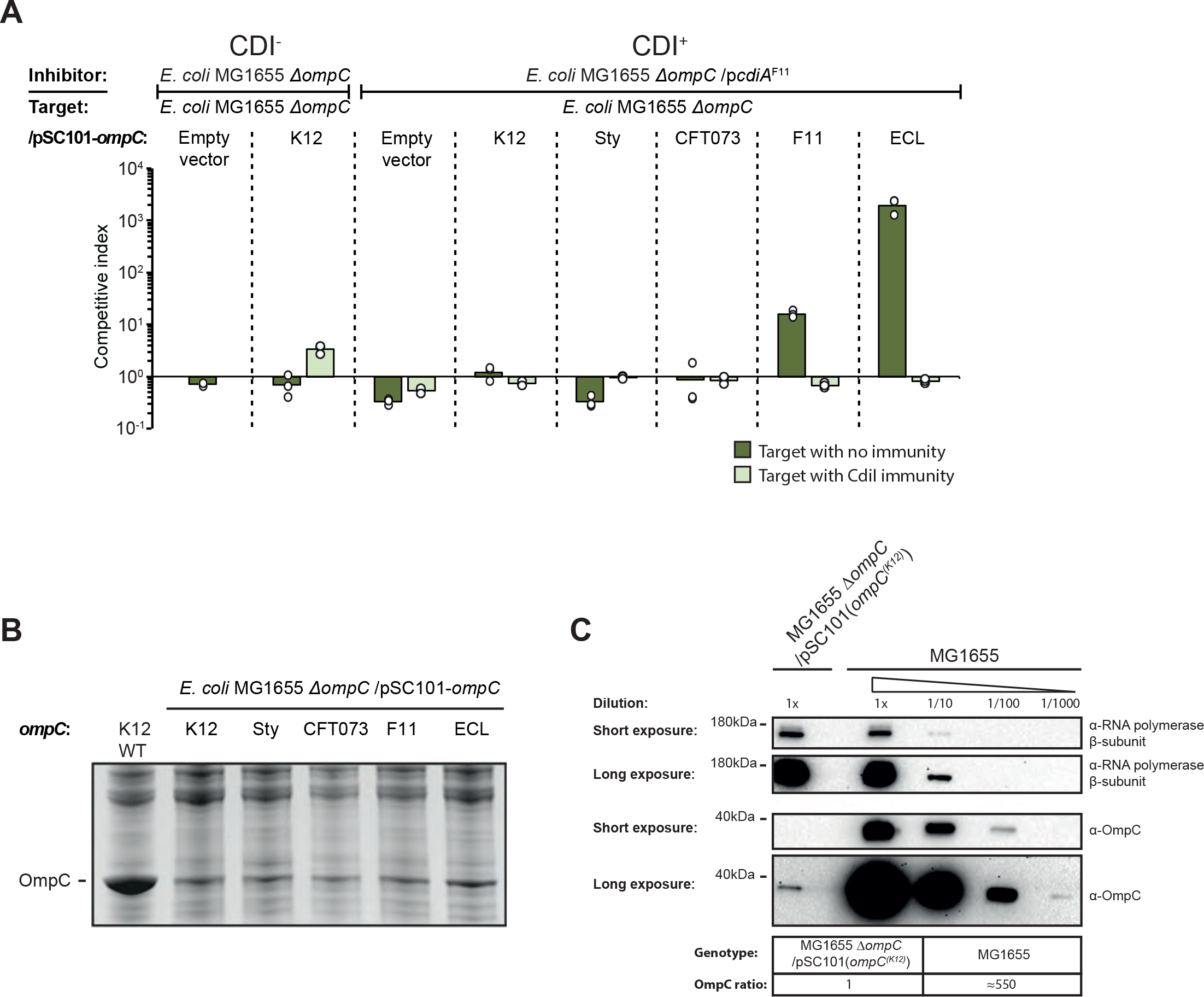
Class II CdiA toxin delivery is affected by OmpC expression levels. **A)** Average competitive index of cells expressing CdiA^F11^ after co-culturing with MG1655 cells expressing different OmpC’s from a low-copy (pSC101) plasmid with (light green bars) or without (dark green bars) CdiI expressed from plasmid (n=3 biological replicates). Cells were co-cultured for 5h in liquid LB media. Individual data points of the biological replicates are shown as black and white circles. **B)** MG1655 cells expressing different OmpC’s from Fig. S3A were grown in LB and outer membrane fractions were enriched and separated by SDS-PAGE. **C)** Western blot of MG1655 cells from figure 1A and S3A expressing OmpC^K12^ either from the chromosome or from the low-copy (pSC101) plasmid.

**Figure S4.**
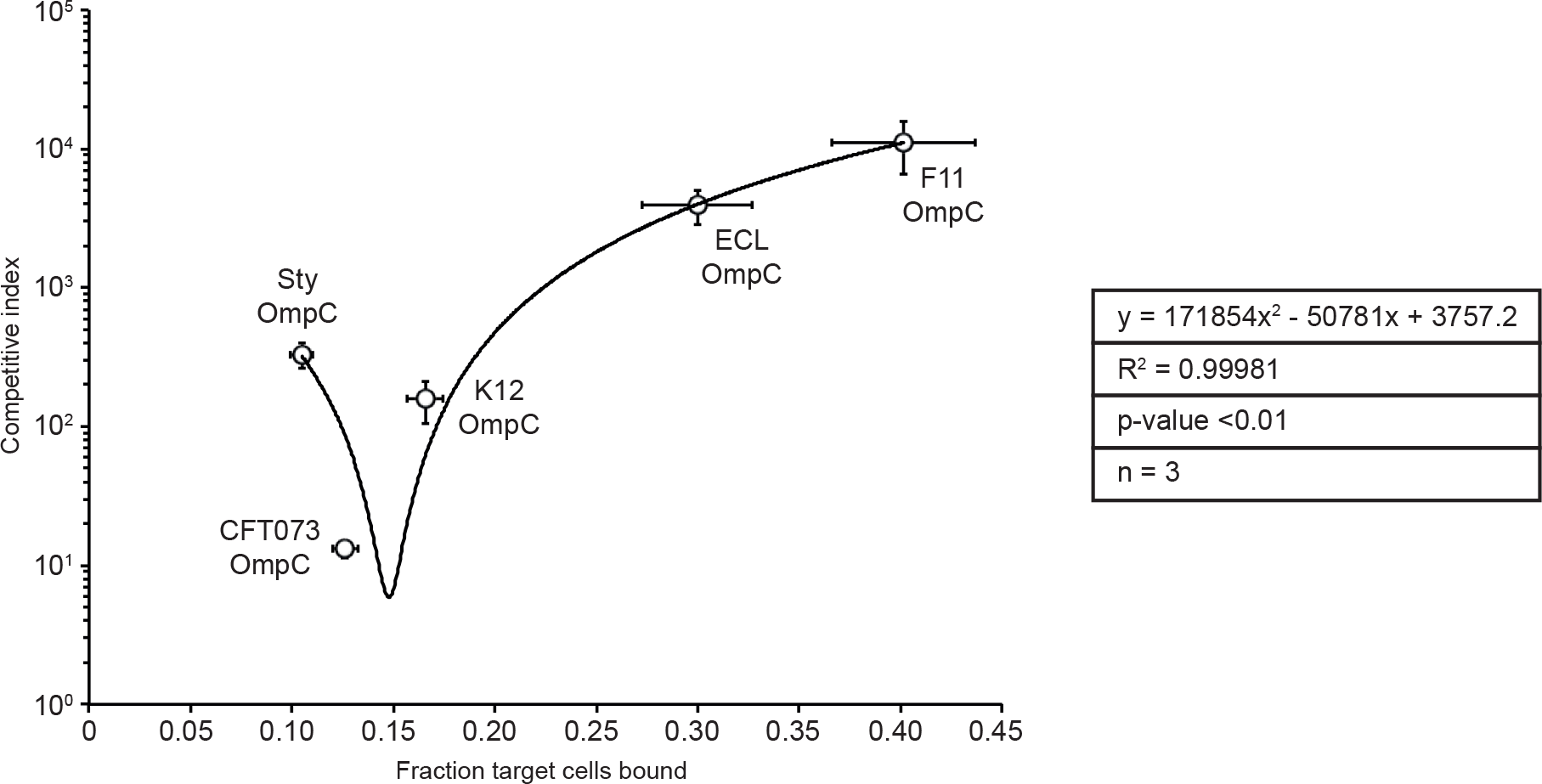
Correlation of cell-cell binding with growth inhibition. Fraction of target cells, expressing different OmpC’s, bound to inhibitor cells expressing CdiA^F11^ was correlated against the measured competitive index of the same strains when co-cultured in LB (n=3 biological replicates). Statistical significance was determined using Pearson correlation tables for R^2^ values.

**Figure S5.**
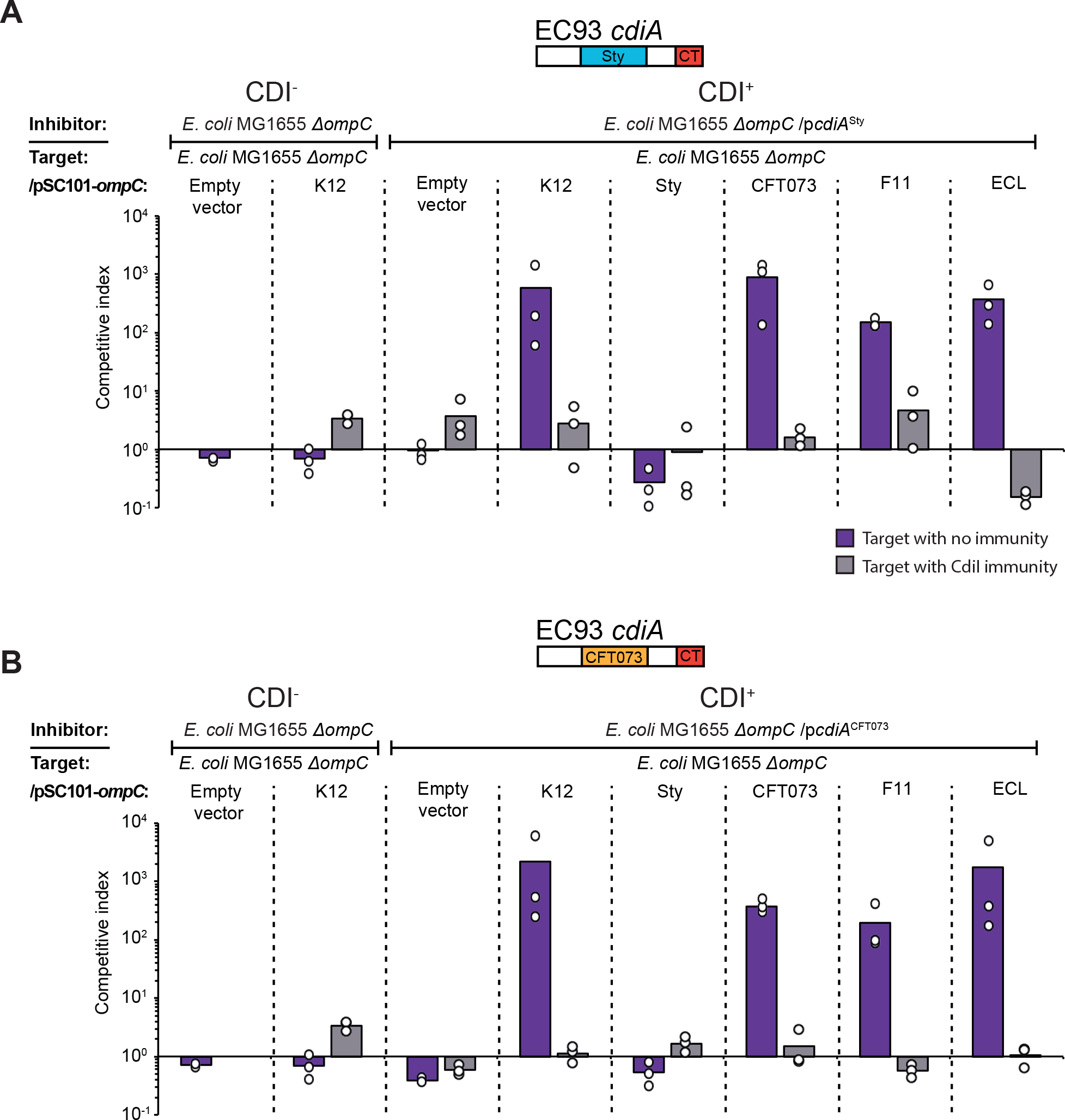
Other Class II CdiA RBD is able to deliver effectors to cells expressing the OmpC receptor from other strains and species on solid media. **A & B)** Average competitive index of cells expressing CdiA^Sty^ (**A**) or CdiA^CFT073^ (**B**) after co-culturing with MG1655 cells expressing different OmpC’s from a low-copy (pSC101) plasmid with (light grey bars) or without (purple bars) CdiI expressed from plasmid (n=3 biological replicates). Cells were co-cultured for 24h on solid M9Glu media. Individual data points of the biological replicates are shown as black and white circles.

**Figure S6.**
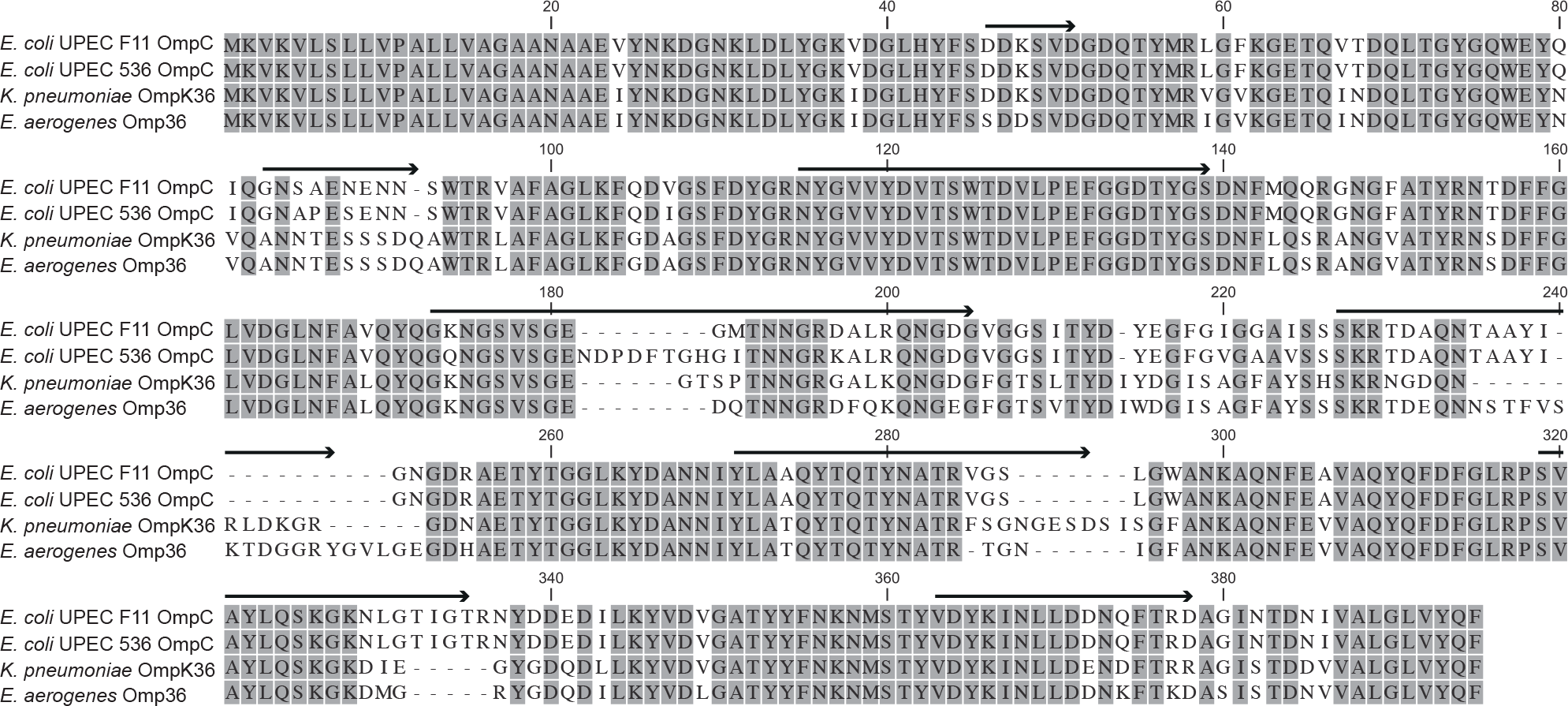
Alignment of OmpC homologs from *Klebsiella pneumoniae* and *Enterobacter aerogenes*. Homologous residues are shown in grey and non-homologous residues are white.

**Figure S7.**
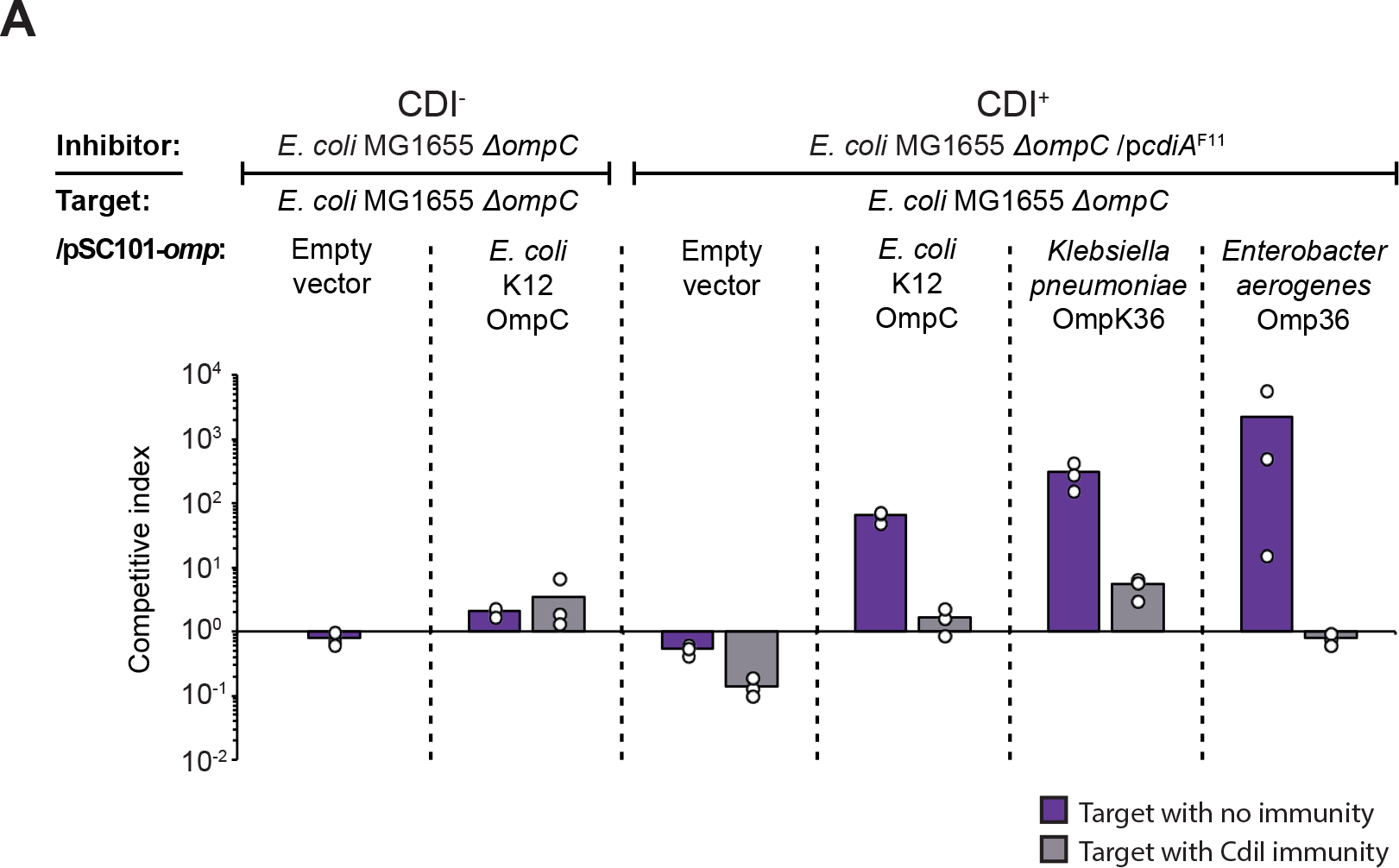
Expression of OmpC homologs from *Klebsiella pneumoniae* and *Enterobacter aerogenes* allow effector delivery into MG1655 cells. Average competitive index of cells expressing CdiA^F11^ (after co-culturing with MG1655 cells expressing different OmpC’s from a low-copy (pSC101) plasmid with (light grey bars) or without (purple bars) CdiI expressed from plasmid (n=3 biological replicates). Cells were co-cultured for 24h on solid M9Glu media. Individual data points of the biological replicates are shown as black and white circles.

**Figure S8.**
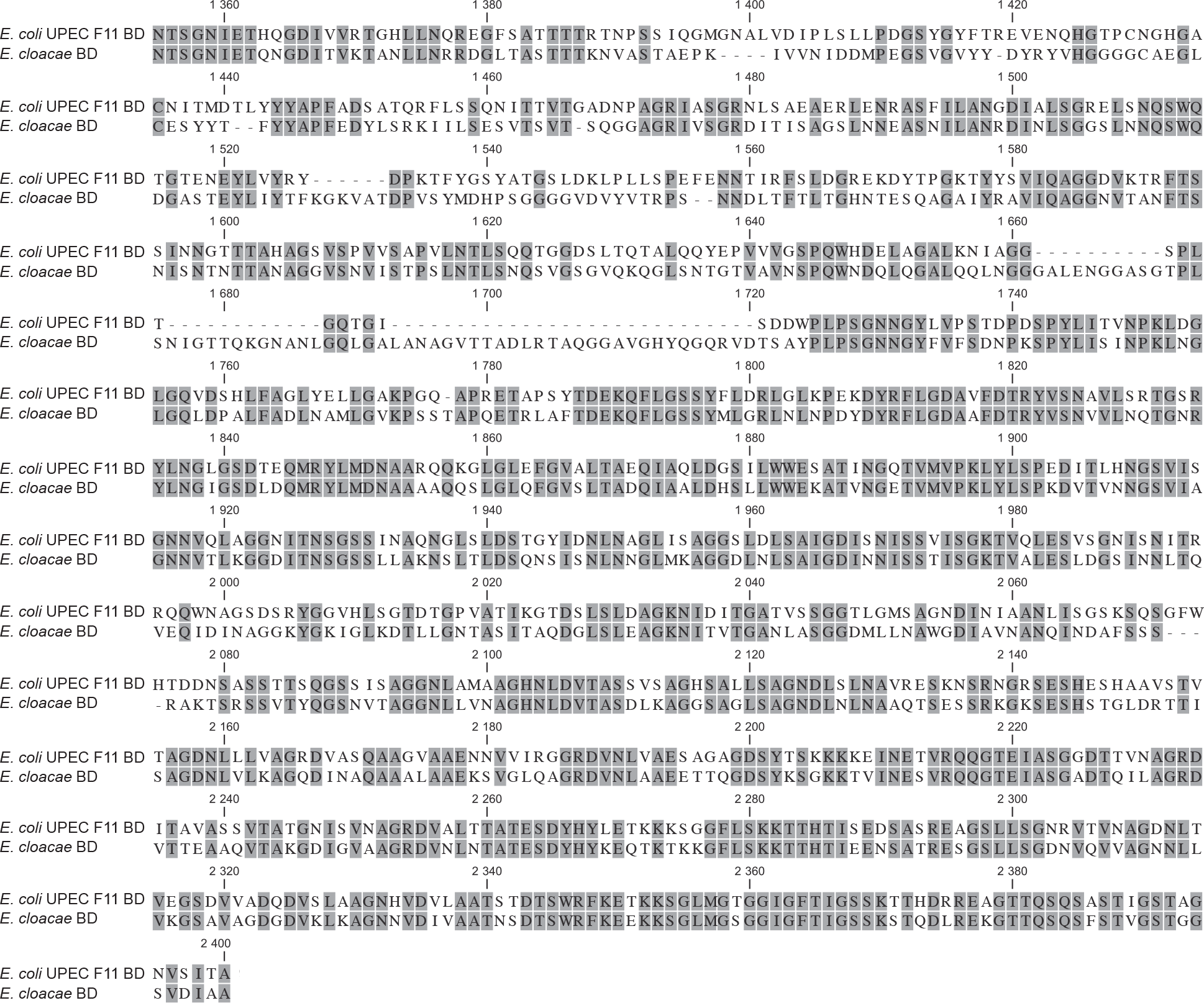
Alignment of CdiA receptor-binding domains from *Escherichia coli* UPEC F11 and *Enterobacter cloacae*. Homologous residues are shown in grey and non-homologous residues are white.

**Figure S9.**
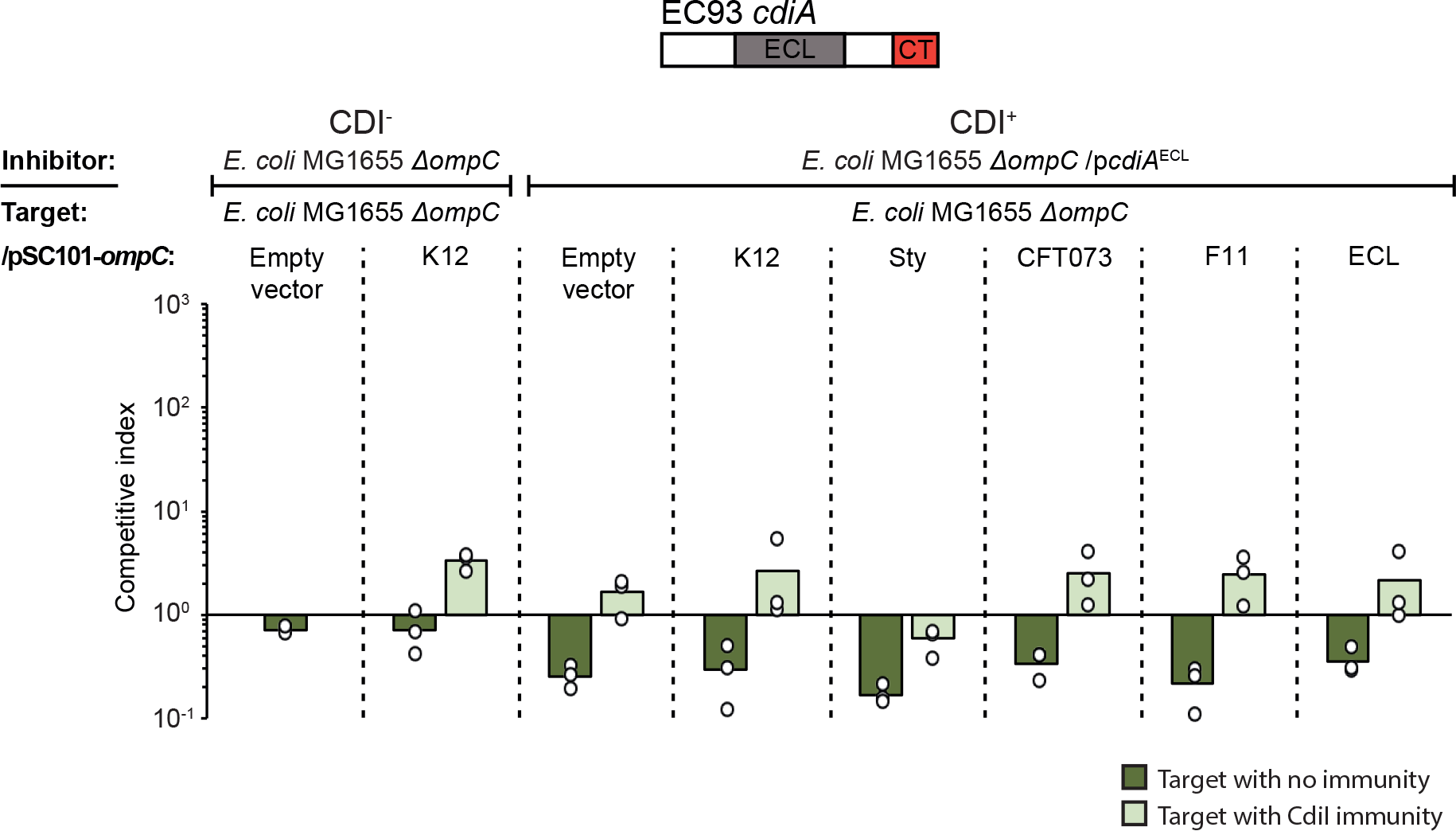
The CdiA protein of E. cloacae does not inhibit *E. coli* MG1655 expressing different OmpC’s in liquid media. Average competitive index of cells expressing CdiA^ECL^ after co-culturing with MG1655 cells expressing different OmpC’s from a constitutive PJ23101 promoter on a low-copy (pSC101) plasmid. Co-culturing was for 5h in liquid LB media with (light green bars) or without (green bars) CdiI expressed from plasmid (n=3 biological replicates). Individual data points of the biological replicates are shown as black and white circles.

## Supplementary methods

### Chromosomal and plasmid constructs

#### Replacement of ompC CDS on the chromosome

A *cat*-*sacB*-*amilCP* selection, counter-selection and screening marker, described previously (Nasvall, 2017), was amplified with primers 1190 and 1191 to introduce 40bp overlap at both 5’ and 3’ ends with homology to the 5’ and 3’ ends of the *ompC* CDS of MG1655. The *cat*-*sacB*-*amilCP* cassette was then integrated onto the genome of MG1655 by lambda red recombination (Datsenko & Wanner, 2000), simultaneously knocking-out *ompC*^(MG1655)^ and generating strain SK2754. *ompC* CDS from *S. typhimurium* LT2, *E. coli* Nissle 1917, *E. coli* F11 and *E. cloacae* ATCC 13047 was amplified with primers 1192 and 1193 and the subsequent PCR products was integrated on the chromosome by lambda red recombination (Datsenko & Wanner, 2000) knocking-out the *cat*-*sacB*-*amilCP* cassette and generating seamless *ompC* CDS swaps (SK2777-SK2780). Positive clones were selected for sucrose tolerance and screened for the lack of blue coloring (result of chromo protein AmilCP expression) and chloramphenicol sensitivity. The constructs were verified by PCR using primers 631 and 632, as well as primers 634 and 635, and further verified by sequencing of the PCR products generated from primers 631 and 632.

#### Construction of pSC101::PJ23101-ompC expressing plasmids

*ompC* was amplified from *S. typhimurium* LT2, *E. coli* K12 MG1655, *E. coli* Nissle 1917, *E. coli* F11 and *E. cloacae* ATCC 13047 using oligos 606-612 as indicated in Table S3 and inserted into the pSC101 plasmid under the control of the PJ23101 promoter. The resulting plasmids were verified by sequencing using oligos 482 and 483.

#### Construction of pCloDF1::PJ23101-cdiI^EC93main^ expression plasmid

*cdiI* was amplified from EC93 using oligos 964 and972 as indicated in Table S3 and cloned into the pCloDF1 plasmid under the control of the PJ23101 promoter. The resulting plasmids were verified by sequencing using oligos 986 and 987.

#### Replacement of the cdiA receptor-binding domains on the medium-copy pcolE1-cdiBAI plasmid

To change the receptor-binding domain of *cdiA*^EC93^, a *cat-sacB* cassette (Nasvall, 2017) with homology upstream and downstream of the receptor-binding domain of *cdiA*^EC93^ was amplified using oligos 583 and 613. The insert was designed to create SrfIrestriction sites upstream and downstream of the receptor-binding domain when inserted. The receptor-binding domains from *E. cloacae* ATCC 13047 and *E. coli* CFT073 *cdiA* was amplified using oligos 794, 795, 584 and 585 and cloned into pDAL660 *cdiA::cat-sacB* using SrfIrestriction sites. A potential class II CDI system from *S.typhi* was identified bioinformatically and the receptor-binding domain was synthesized (Gene art, Thermo Scientific, USA) and cloned as described for the other constructs. The resulting plasmids were transformed into *E. coli* K12 MG1655 *ompC::kan* and positive clones were selected for sucrose tolerance and screened for Cam sensitivity, followed by PCR verification and sequencing with primers indicated in table S2. The resulting plasmids carrying the *cdiBAI*^EC93^ system with the receptor binding domain swaps was transformed into *E. coli* K12 MG1655 *ompC::kan*.

#### sYFP2 reporter plasmid

The *pJ23101-sYFP2* and *osmY-mtagBFP* reporter plasmid (SK2292), previously described in (Ghosh et al., 2018) was modified to remove the ClpXP dependent degradation tags fused to both sYFP2 and mtagBFP. The pSC101 plasmid was amplified with primers 1062 and 1276, the resulting PCR-products were gel-purified followed by re-circularized with T4 DNA ligase (Thermo Scientific, USA) and transformation into NEB5alfa (New England Biolabs, USA) generating plasmid SK2876. Positive clones were verified by sequencing using the same primers described before (Ghosh et al., 2018). The SK2876 plasmid was amplified with primers 1050 and 1266, the resulting PCR-products were then gel-purified followed by re-circularized with T4 DNA ligase (Thermo Scientific, USA) and transformation into NEB5alfa (New England Biolabs, USA) generating plasmid SK2938. Positive clones were verified same as above.

**Table S1.**
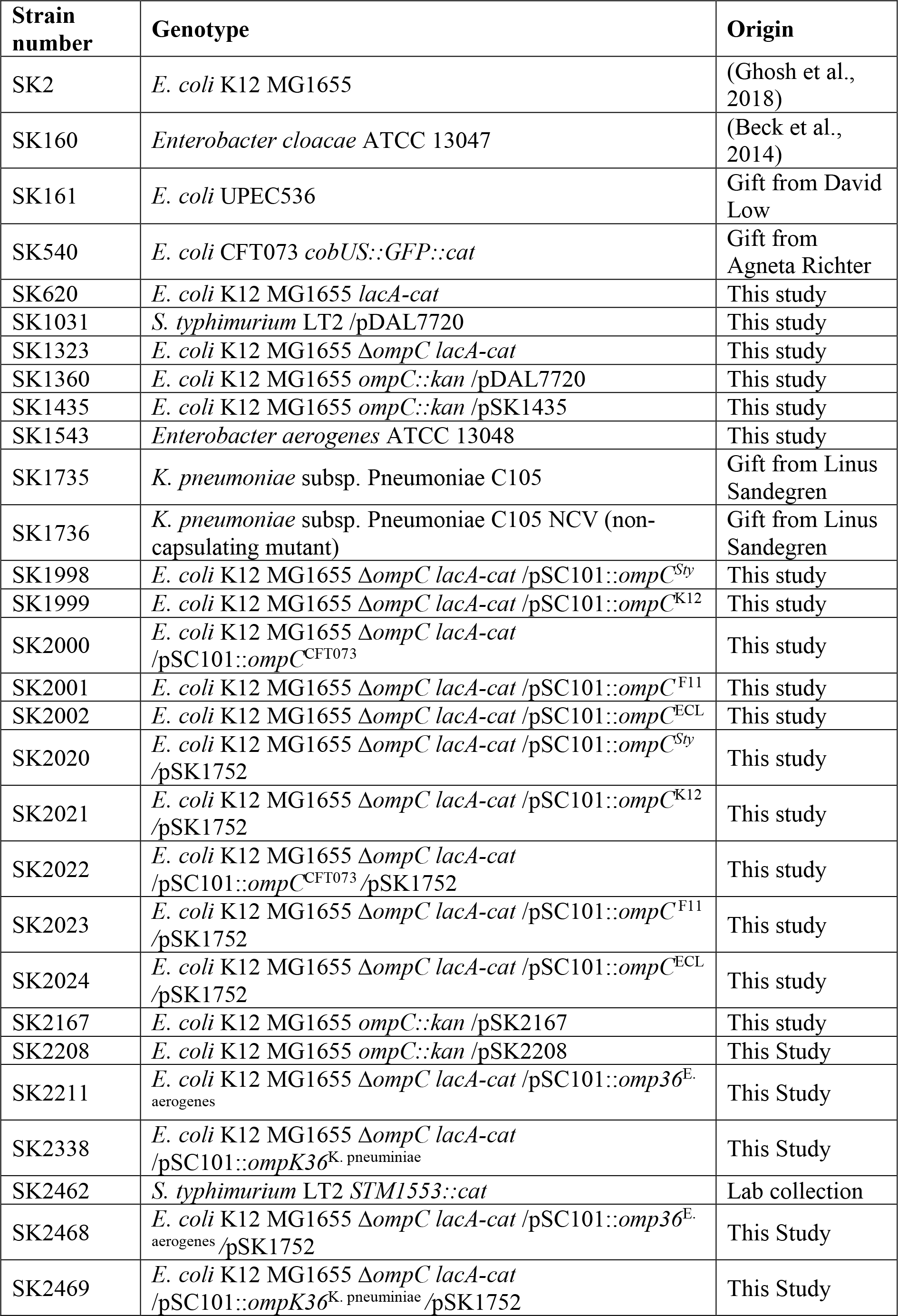
Bacterial strains used in this study.

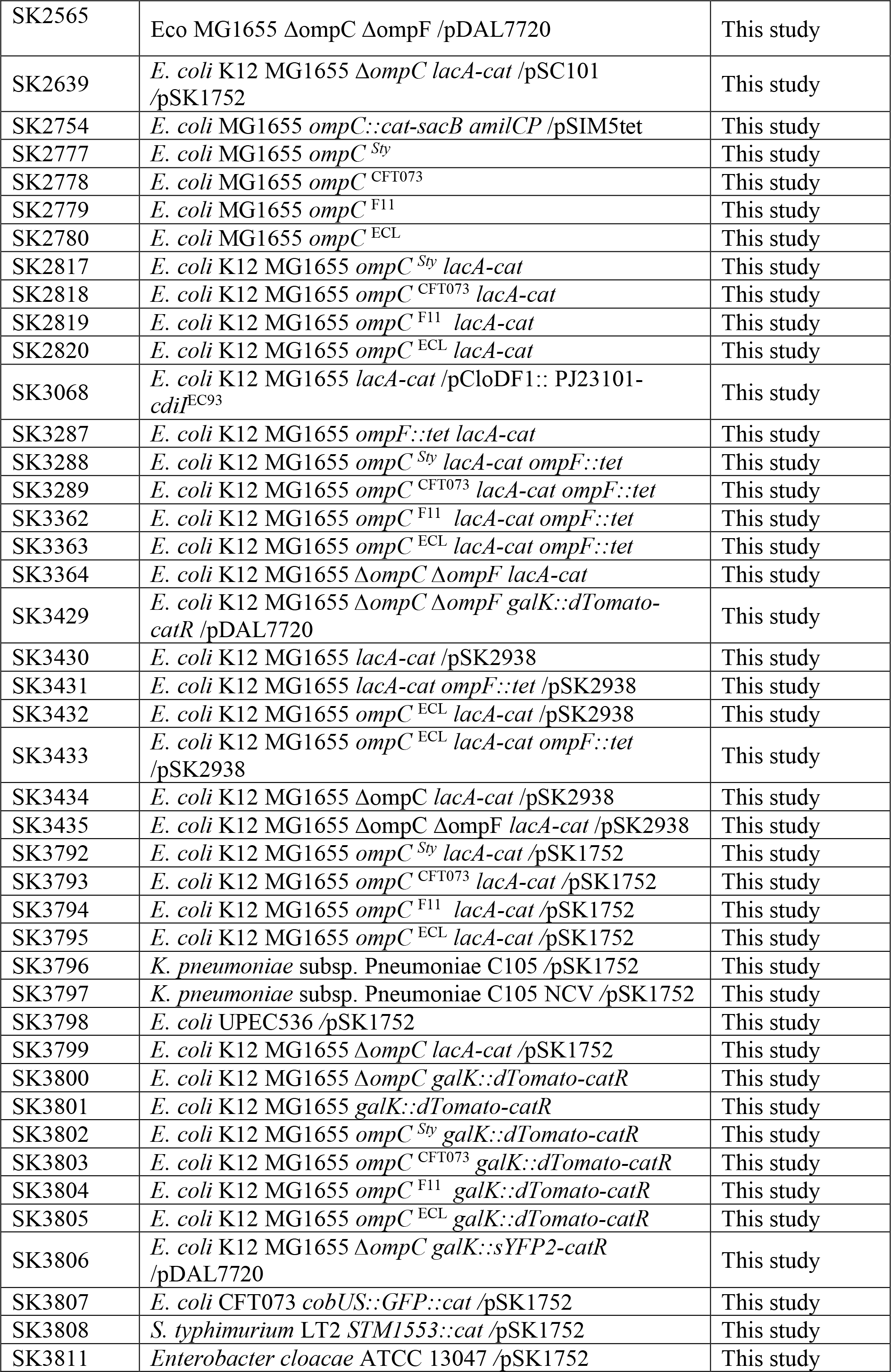

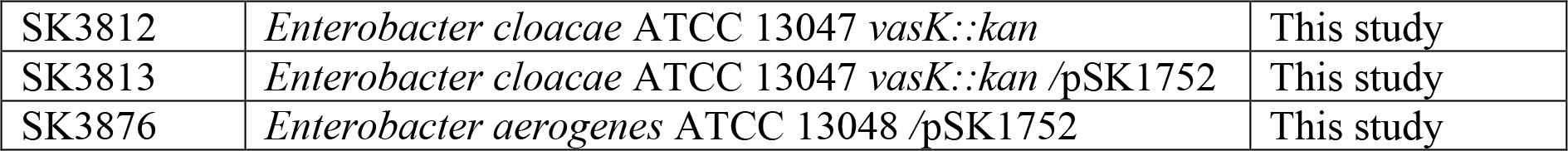

**Table S2.**
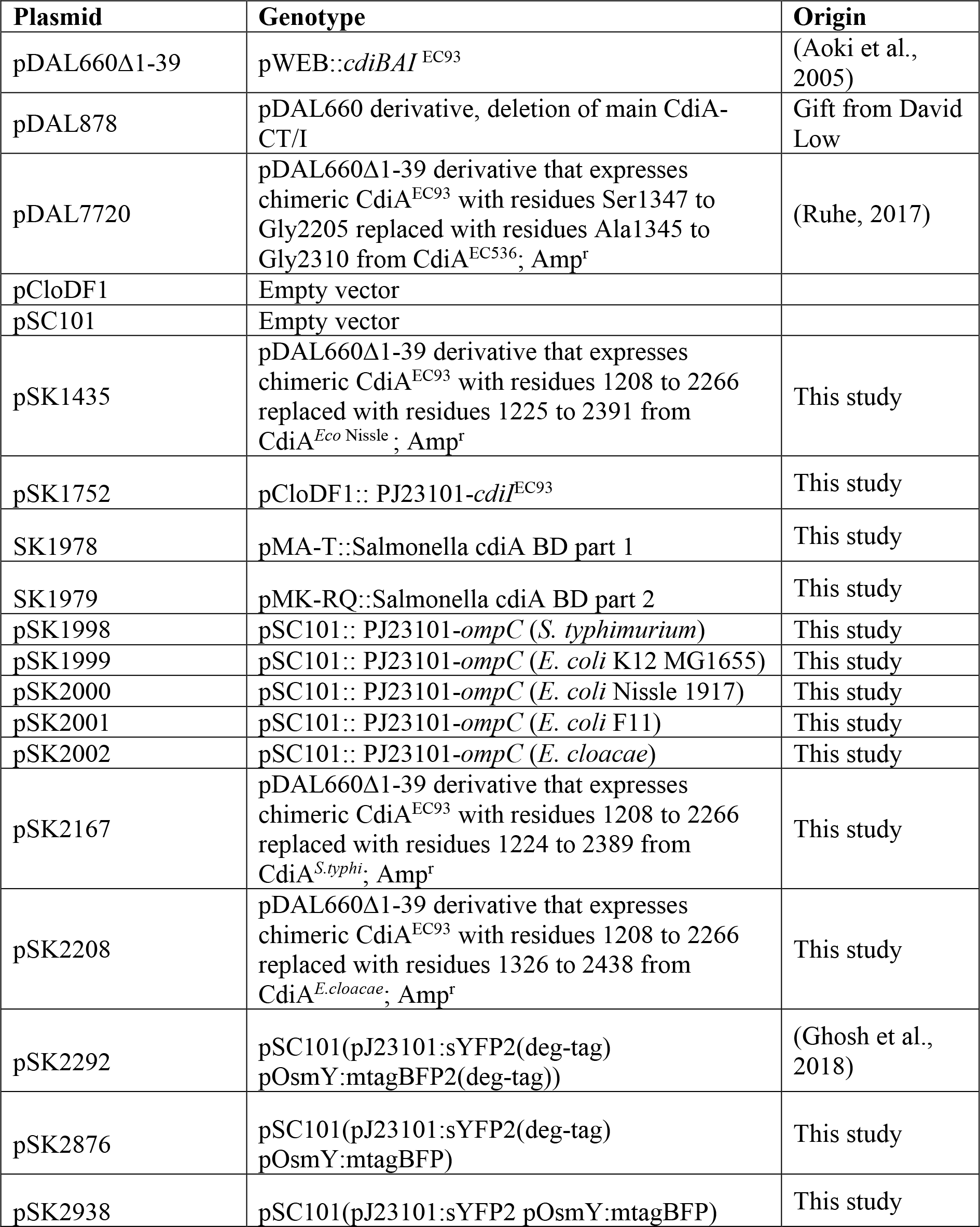
Plasmids used in this study.

| Plasmid | Genotype | Origin |
| --- | --- | --- |
| pDAL660Δ1-39 | pWEB::*cdiBAI*<sup>EC93</sup> | (Aoki et al., 2005) |
| pDAL878 | pDAL660 derivative, deletion of main CdiA-CT/I | Gift from David Low |
| pDAL7720 | pDAL660Δ1-39 derivative that expresses chimeric CdiA<sup>EC93</sup> with residues Ser1347 to Gly2205 replaced with residues Ala1345 to Gly2310 from CdiA<sup>EC536</sup>; Amp<sup>r</sup> | (Ruhe, 2017) |
| pCloDF1 | Empty vector |  |
| pSC101 | Empty vector |  |
| pSK1435 | pDAL660Δ1-39 derivative that expresses chimeric CdiA<sup>EC93</sup> with residues 1208 to 2266 replaced with residues 1225 to 2391 from CdiA<sup>Eco Nissle</sup>; Amp<sup>r</sup> | This study |
| pSK1752 | pCloDF1:: PJ23101-*cdiI*<sup>EC93</sup> | This study |
| SK1978 | pMA-T::Salmonella *cdiA* BD part 1 | This study |
| SK1979 | pMK-RQ::Salmonella *cdiA* BD part 2 | This study |
| pSK1998 | pSC101:: PJ23101-*ompC* (*S. typhimurium*) | This study |
| pSK1999 | pSC101:: PJ23101-*ompC* (*E. coli* K12 MG1655) | This study |
| pSK2000 | pSC101:: PJ23101-*ompC* (*E. coli* Nissle 1917) | This study |
| pSK2001 | pSC101:: PJ23101-*ompC* (*E. coli* F11) | This study |
| pSK2002 | pSC101:: PJ23101-*ompC* (*E. cloacae*) | This study |
| pSK2167 | pDAL660Δ1-39 derivative that expresses chimeric CdiA<sup>EC93</sup> with residues 1208 to 2266 replaced with residues 1224 to 2389 from CdiA<sup>S.typhi</sup>; Amp<sup>r</sup> | This study |
| pSK2208 | pDAL660Δ1-39 derivative that expresses chimeric CdiA<sup>EC93</sup> with residues 1208 to 2266 replaced with residues 1326 to 2438 from CdiA<sup>E.cloacae</sup>; Amp<sup>r</sup> | This study |
| pSK2292 | pSC101(pJ23101:sYFP2(deg-tag)<br>pOsmY:mtagBFP2(deg-tag)) | (Ghosh et al., 2018) |
| pSK2876 | pSC101(pJ23101:sYFP2(deg-tag)<br>pOsmY:mtagBFP) | This study |
| pSK2938 | pSC101(pJ23101:sYFP2 pOsmY:mtagBFP) | This study |

**Table S3.**
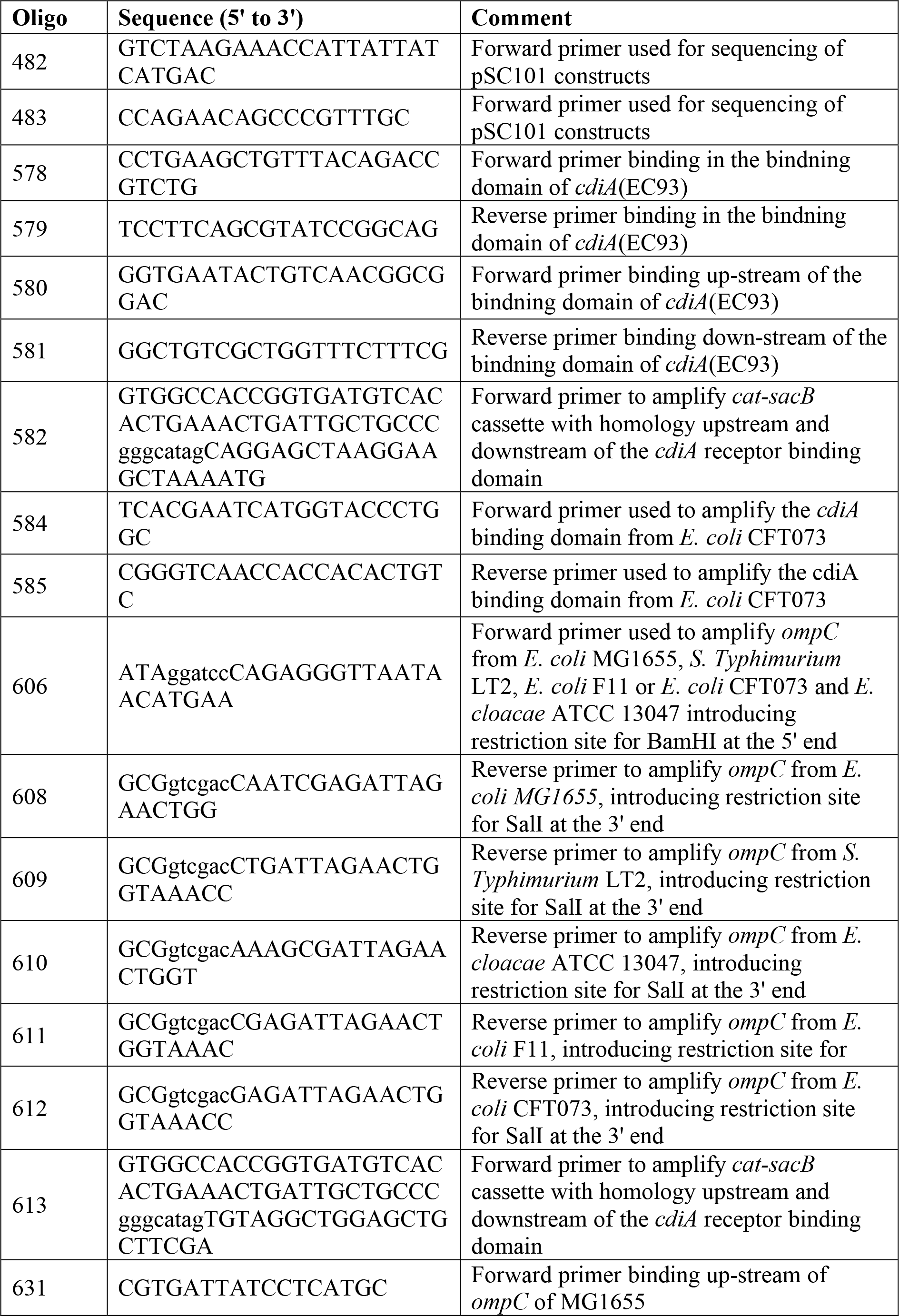
Oligos used in this study.

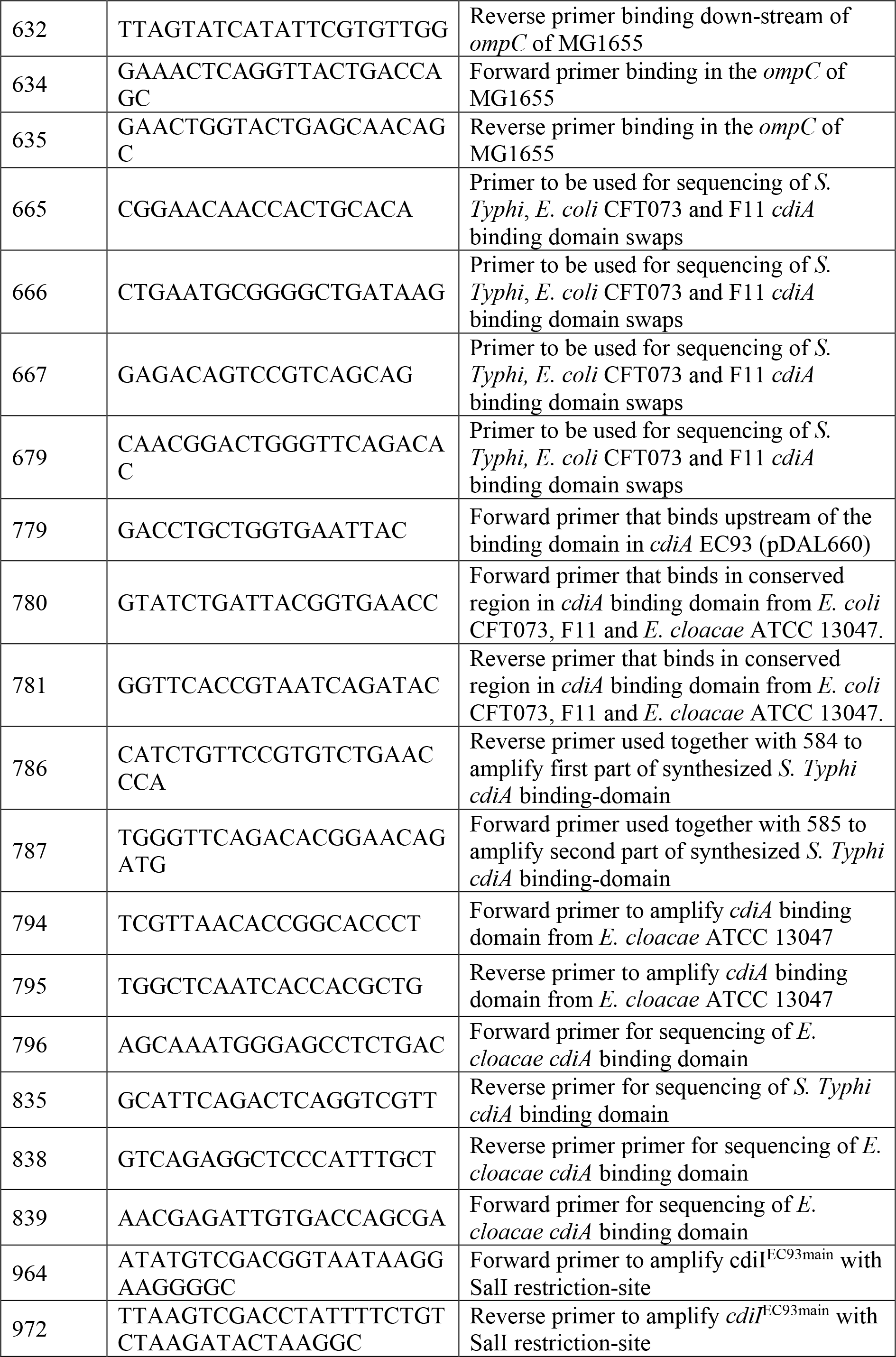

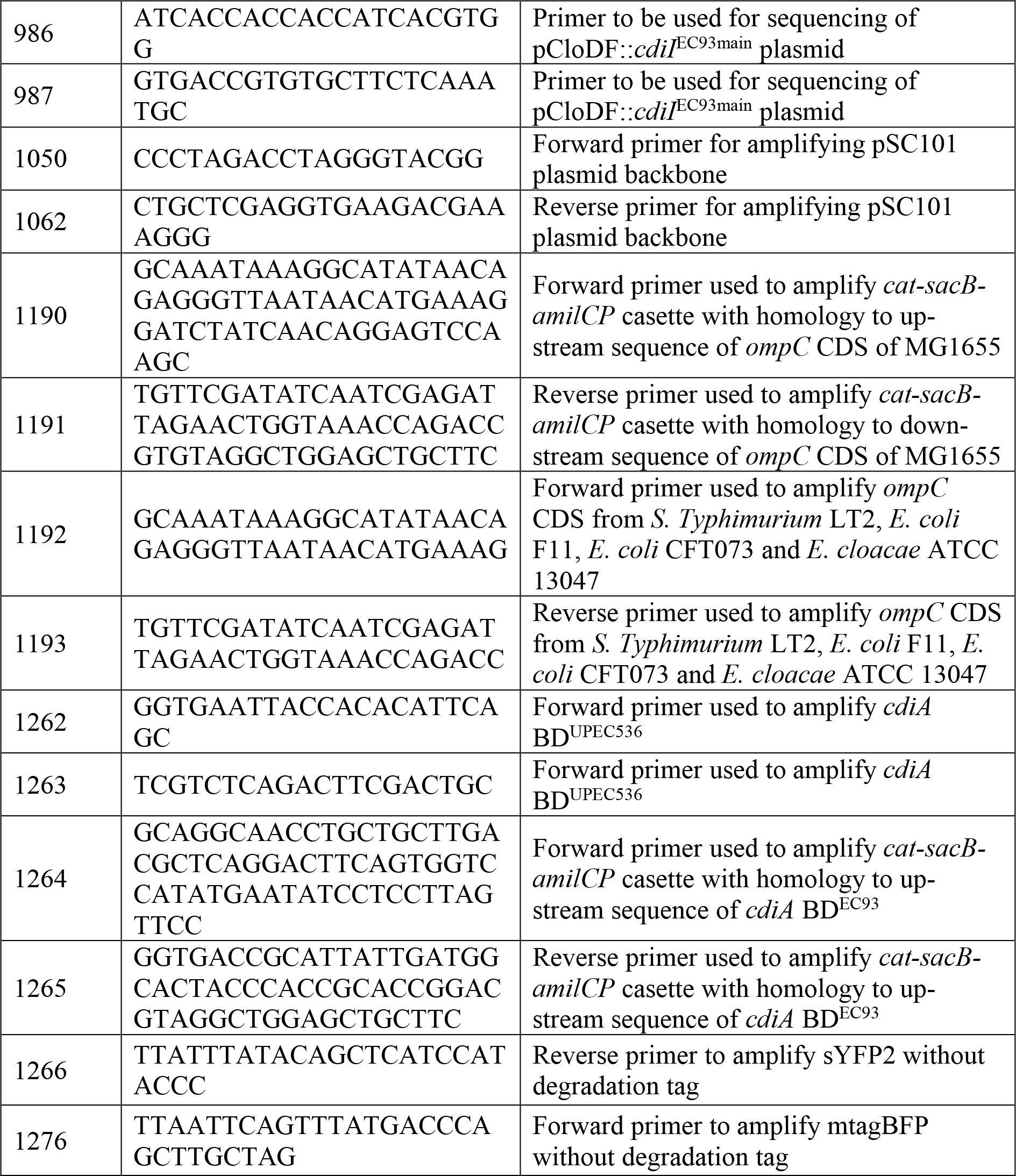

## References

Aoki, S. K., Diner, E. J., de Roodenbeke, C. T., Burgess, B. R., Poole, S. J., Braaten, B. A., … Low, D. A. (2010). A widespread family of polymorphic contact-dependent toxin delivery systems in bacteria. Nature, 468(7322), 439–442. doi:10.1038/nature09490

Aoki, S. K., Malinverni, J. C., Jacoby, K., Thomas, B., Pamma, R., Trinh, B. N., … Low, D. A. (2008). Contact-dependent growth inhibition requires the essential outer membrane protein BamA (YaeT) as the receptor and the inner membrane transport protein AcrB. Molecular microbiology, 70(2), 323–340. doi:10.1111/j.1365-2958.2008.06404.x

Aoki, S. K., Pamma, R., Hernday, A. D., Bickham, J. E., Braaten, B. A., & Low, D. A. (2005). Contact-dependent inhibition of growth in Escherichia coli. Science, 309(5738), 1245–1248. doi:10.1126/science.1115109

Aoki, S. K., Webb, J. S., Braaten, B. A., & Low, D. A. (2009). Contact-dependent growth inhibition causes reversible metabolic downregulation in Escherichia coli. J Bacteriol, 191(6), 1777–1786. doi:10.1128/JB.01437-08

Baba, T., Ara, T., Hasegawa, M., Takai, Y., Okumura, Y., Baba, M., … Mori, H. (2006). Construction of Escherichia coli K-12 in-frame, single-gene knockout mutants: the Keio collection. Mol Syst Biol, 2, 2006 0008. doi:10.1038/msb4100050

Beck, C. M., Morse, R. P., Cunningham, D. A., Iniguez, A., Low, D. A., Goulding, C. W., & Hayes, C. S. (2014). CdiA from Enterobacter cloacae delivers a toxic ribosomal RNase into target bacteria. Structure, 22(5), 707–718. doi:10.1016/j.str.2014.02.012

Beck, C. M., Willett, J. L., Cunningham, D. A., Kim, J. J., Low, D. A., & Hayes, C. S. (2016). CdiA Effectors from Uropathogenic Escherichia coli Use Heterotrimeric Osmoporins as Receptors to Recognize Target Bacteria. PLoS Pathog, 12(10), e1005925. doi:10.1371/journal.ppat.1005925

Bertozzi Silva, J., Storms, Z., & Sauvageau, D. (2016). Host receptors for bacteriophage adsorption. FEMS Microbiol Lett, 363(4). doi:10.1093/femsle/fnw002

Cao, P., & Wall, D. (2017). Self-identity reprogrammed by a single residue switch in a cell surface receptor of a social bacterium. Proceedings of the National Academy of Sciences of the United States of America, 114(14), 3732–3737. doi:10.1073/pnas.1700315114

Cao, Z., Casabona, M. G., Kneuper, H., Chalmers, J. D., & Palmer, T. (2016). The type VII secretion system of Staphylococcus aureus secretes a nuclease toxin that targets competitor bacteria. Nat Microbiol, 2, 16183. doi:10.1038/nmicrobiol.2016.183

Forst, S., Delgado, J., Ramakrishnan, G., & Inouye, M. (1988). Regulation of ompC and ompF expression in Escherichia coli in the absence of envZ. J Bacteriol, 170(11), 5080–5085.

Garcia, E. C., Perault, A. I., Marlatt, S. A., & Cotter, P. A. (2016). Interbacterial signaling via Burkholderia contact-dependent growth inhibition system proteins. Proc Natl Acad Sci U S A, 113(29), 8296–8301. doi:10.1073/pnas.1606323113

Garcia-Bayona, L., Guo, M. S., & Laub, M. T. (2017). Contact-dependent killing by Caulobacter crescentus via cell surface-associated, glycine zipper proteins. Elife, 6. doi:10.7554/eLife.24869

Ghosh, A., Baltekin, O., Waneskog, M., Elkhalifa, D., Hammarlof, D. L., Elf, J., & Koskiniemi, S. (2018). Contact-dependent growth inhibition induces high levels of antibiotic-tolerant persister cells in clonal bacterial populations. EMBO J. doi:10.15252/embj.201798026

Gibbs, K. A., Urbanowski, M. L., & Greenberg, E. P. (2008). Genetic determinants of self identity and social recognition in bacteria. Science, 321(5886), 256–259. doi:10.1126/science.1160033

Hayes, C. S., Aoki, S. K., & Low, D. A. (2010). Bacterial contact-dependent delivery systems. Annu Rev Genet, 44, 71–90. doi:10.1146/annurev.genet.42.110807.091449

Hood, R. D., Singh, P., Hsu, F., Guvener, T., Carl, M. A., Trinidad, R. R., … Mougous, J. D. (2010). A type VI secretion system of Pseudomonas aeruginosa targets a toxin to bacteria. Cell Host Microbe, 7(1), 25–37. doi:10.1016/j.chom.2009.12.007

Kelly, J. R., Rubin, A. J., Davis, J. H., Ajo-Franklin, C. M., Cumbers, J., Czar, M. J., … Endy, D. (2009). Measuring the activity of BioBrick promoters using an in vivo reference standard. J Biol Eng, 3, 4. doi:10.1186/1754-1611-3-4

Koskiniemi, S., Garza-Sanchez, F., Edman, N., Chaudhuri, S., Poole, S. J., Manoil, C., … Low, D. A. (2015). Genetic analysis of the CDI pathway from Burkholderia pseudomallei 1026b. PLoS One, 10(3), e0120265. doi:10.1371/journal.pone.0120265

Leo, J. C., Grin, I., & Linke, D. (2012). Type V secretion: mechanism(s) of autotransport through the bacterial outer membrane. Philosophical Transactions of the Royal Society B-Biological Sciences, 367(1592), 1088–1101. doi:10.1098/rstb.2011.0208

Li, G. W., Burkhardt, D., Gross, C., & Weissman, J. S. (2014). Quantifying absolute protein synthesis rates reveals principles underlying allocation of cellular resources. Cell, 157(3), 624–635. doi:10.1016/j.cell.2014.02.033

Liu, Y. F., Yan, J. J., Lei, H. Y., Teng, C. H., Wang, M. C., Tseng, C. C., & Wu, J. J. (2012). Loss of outer membrane protein C in Escherichia coli contributes to both antibiotic resistance and escaping antibody-dependent bactericidal activity. Infect Immun, 80(5), 1815–1822. doi:10.1128/IAI.06395-11

Morse, R. P., Nikolakakis, K. C., Willett, J. L., Gerrick, E., Low, D. A., Hayes, C. S., & Goulding, C. W. (2012). Structural basis of toxicity and immunity in contact-dependent growth inhibition (CDI) systems. Proc Natl Acad Sci U S A, 109(52), 21480–21485. doi:10.1073/pnas.1216238110

Pathak, D. T., Wei, X., Dey, A., & Wall, D. (2013). Molecular recognition by a polymorphic cell surface receptor governs cooperative behaviors in bacteria. PLoS genetics, 9(11), e1003891. doi:10.1371/journal.pgen.1003891

Ruhe, Z. C., Low, D. A., & Hayes, C. S. (2013). Bacterial contact-dependent growth inhibition. Trends in Microbiology, 21(5), 230–237. doi:10.1016/j.tim.2013.02.003

Ruhe, Z. C., Nguyen, J. Y., Xiong, J., Koskiniemi, S., Beck, C. M., Perkins, B. R., … Hayes, C. S. (2017). CdiA Effectors Use Modular Receptor-Binding Domains To Recognize Target Bacteria. MBio, 8(2). doi:10.1128/mBio.00290-17

Ruhe, Z. C., Subramanian, P., Song, K., Nguyen, J. Y., Stevens, T. A., Low, D. A., … Hayes, C. S. (2018). Programmed Secretion Arrest and Receptor-Triggered Toxin Export during Antibacterial Contact-Dependent Growth Inhibition. Cell, 175(4), 921–933 e914. doi:10.1016/j.cell.2018.10.033

Ruhe, Z. C., Townsley, L., Wallace, A. B., King, A., Van der Woude, M. W., Low, D. A., … Hayes, C. S. (2015). CdiA promotes receptor-independent intercellular adhesion. Mol Microbiol, 98(1), 175–192. doi:10.1111/mmi.13114

Ruhe, Z. C., Wallace, A. B., Low, D. A., & Hayes, C. S. (2013). Receptor polymorphism restricts contact-dependent growth inhibition to members of the same species. MBio, 4(4). doi:10.1128/mBio.00480-13

Russell, A. B., Peterson, S. B., & Mougous, J. D. (2014). Type VI secretion system effectors: poisons with a purpose. Nat Rev Microbiol, 12(2), 137–148. doi:10.1038/nrmicro3185

Schuman, W. (2006). Dynamics of the Bacterial Chromosome: Structure and Function: Wiley-VCH Verlag GmbH &Co. KGaA.

Singh, S. P., Williams, Y. U., Klebba, P. E., Macchia, P., & Miller, S. (2000). Immune recognition of porin and lipopolysaccharide epitopes of Salmonella typhimurium in mice. Microb Pathog, 28(3), 157–167. doi:10.1006/mpat.1999.0332

Souza, D. P., Oka, G. U., Alvarez-Martinez, C. E., Bisson-Filho, A. W., Dunger, G., Hobeika, L., … Farah, C. S. (2015). Bacterial killing via a type IV secretion system. Nat Commun, 6, 6453. doi:10.1038/ncomms7453

Stenkova, A. M., Bystritskaya, E. P., Guzev, K. V., Rakin, A. V., & Isaeva, M. P. (2016). Molecular Evolution of the Yersinia Major Outer Membrane Protein C (OmpC). Evol Bioinform Online, 12, 185–191. doi:10.4137/EBO.S40346

Vassallo, C. N., Cao, P., Conklin, A., Finkelstein, H., Hayes, C. S., & Wall, D. (2017). Infectious polymorphic toxins delivered by outer membrane exchange discriminate kin in myxobacteria. Elife, 6. doi:10.7554/eLife.29397

Wall, D. (2016). Kin Recognition in Bacteria. Annu Rev Microbiol, 70, 143–160. doi:10.1146/annurev-micro-102215-095325

Webb, J. S., Nikolakakis, K. C., Willett, J. L., Aoki, S. K., Hayes, C. S., & Low, D. A. (2013). Delivery of CdiA nuclease toxins into target cells during contact-dependent growth inhibition. PloS one, 8(2), e57609. doi:10.1371/journal.pone.0057609

Wenren, L. M., Sullivan, N. L., Cardarelli, L., Septer, A. N., & Gibbs, K. A. (2013). Two independent pathways for self-recognition in Proteus mirabilis are linked by type VI-dependent export. MBio, 4(4). doi:10.1128/mBio.00374-13

Willett, J. L., Gucinski, G. C., Fatherree, J. P., Low, D. A., & Hayes, C. S. (2015). Contact-dependent growth inhibition toxins exploit multiple independent cell-entry pathways. Proc Natl Acad Sci U S A, 112(36), 11341–11346. doi:10.1073/pnas.1512124112

## References

Datsenko, K. A., & Wanner, B. L. (2000). One-step inactivation of chromosomal genes in Escherichia coli K-12 using PCR products. Proceedings of the National Academy of Sciences of the United States of America, 97(12), 6640–6645. doi:10.1073/pnas.120163297

Nasvall, J. (2017). Direct and Inverted Repeat stimulated excision (DIRex): Simple, single-step, and scar-free mutagenesis of bacterial genes. PLoS One, 12(8), e0184126. doi:10.1371/journal.pone.0184126

Ruhe, Z. C., Nguyen, J. Y., Xiao, J., Koskiniemi, S., Beck, C. M., Perkins, B., Low, D. A., Hayes, C. S. (2017). CdiA effectors use modular receptor-binding domains to recognize target bacteria. Submitted manuscript.

